# A Retinoic Acid:YAP1 signaling axis controls atrial lineage commitment

**DOI:** 10.1101/2024.07.11.602981

**Authors:** Elizabeth Abraham, Brett Volmert, Thomas Roule, Ling Huang, Jingting Yu, April E. Williams, Henry M. Cohen, Aidan Douglas, Emily Megill, Alex Morris, Eleonora Stronati, Raquel Fueyo, Mikel Zubillaga, John W. Elrod, Naiara Akizu, Aitor Aguirre, Conchi Estaras

## Abstract

Vitamin A/Retinoic Acid (Vit A/RA) signaling is essential for heart development. In cardiac progenitor cells (CPCs), RA signaling induces the expression of atrial lineage genes while repressing ventricular genes, thereby promoting the acquisition of an atrial cardiomyocyte cell fate. To achieve this, RA coordinates a complex regulatory network of downstream effectors that is not fully identified. To address this gap, we applied a functional genomics approach (i.e scRNAseq and snATACseq) to untreated and RA-treated human embryonic stem cells (hESCs)-derived CPCs. Unbiased analysis revealed that the Hippo effectors YAP1 and TEAD4 are integrated with the atrial transcription factor enhancer network, and that YAP1 is necessary for activation of RA-enhancers in CPCs. Furthermore, in vivo analysis of control and conditionally YAP1 KO mouse embryos (*Sox2-cre*) revealed that the expression of atrial lineage genes, such as *NR2F2*, is compromised by YAP1 deletion in the CPCs of the second heart field. Accordingly, we found that YAP1 is required for the formation of an atrial chamber but is dispensable for the formation of a ventricle, in hESC-derived patterned cardiac organoids. Overall, our findings revealed that YAP1 is a non-canonical effector of RA signaling essential for the acquisition of atrial lineages during cardiogenesis.

## INTRODUCTION

Congenital heart defects (CHD) affect ∼1% of newborns and the etiology is unknown in over 70% of cases. Growing evidence shows that the origins of the majority of CHD are related to pathogenic mechanisms that disturb the specification and differentiation of cardiac progenitor cells (CPC)^1,2^. Vitamin A (Vit A) signaling pathway regulates the differentiation of CPCs. Consequently, Vit A has been linked to atrial, septum and conotruncal defects, when present in excess or deficiency due to mutations or nutritional factors^3–7^. However, infants with normal in utero exposure to Vit A and lack of related mutations still manifest Vit A-associated defects, suggesting that unknown factors regulate Vit A activity and potentially contribute to CHD.

Vitamin A is an essential dietary vitamin. It undergoes metabolic conversion to Retinoic Acid (RA) through a series of enzymatic reactions, with RA acting as a nuclear ligand of RA Receptors (RARs) to regulate gene transcription^8^. In early development, RA signaling regulates the posterior-to-anterior patterning of the CPC pool located in the second heart field (SHF) of the embryo through two important effects: RA signaling **i)** restricts the expression of anterior SHF (AHF) genes such as *Fgf8* and *Isl1* and **ii)** activates genes that promote posterior differentiation of the SHF (pSHF) and acquisition of atrial and sinus venosus lineages, such as *Nr2f2*, *Meis2* or *Hoxa1*^4,5,9–12^. Fine-tuned expression of such RA-target genes is necessary to control the cell-fate decisions of the SHF; when perturbed, it may lead to a variety of CHD, including Tetralogy of Fallot and septum defects^13,14^. Conveniently, the activity of RA can be recapitulated in both hESC-derived CPCs (hCPCs)^15,16^ and hESC-derived cardiac organoids^16^. These human models are excellent platforms to study how RA regulates the gene program that guides the cell-fate decisions of CPCs.

In order for RA to successfully generate an atrial lineage, it induces the expression of *Nr2f2*, an orphan nuclear receptor that regulates transcription of lineage genes. *Nr2f2* is *necessary and sufficient* to confer atrial identity^17–20^ and cardiomyocyte-specific *Nr2f2* ablation produces a *ventricularized* atrial chamber, due to ectopic expression of ventricular genes in the atrial tissue^19^. Moreover, human mutations in *Nr2f2* lead to CHD^21–23^. *Nr2f2* functions directly as both transcriptional activator and repressor. For instance, *Nr2f2* directly binds and activates atrial genes, such as *TBX5*, while it represses typical ventricular genes *Lbh*, *Irx4* and *Hey2*^19^. With only few *Nr2f2* functional interactors identified to date^24,25^, the factors regulating *Nr2f2* levels and/or its transcriptional output remain elusive.

Furthermore, the RA mechanisms must be precisely regulated in each cellular context as RA elicits distinct responses in diverse cell types during developmental processes. It is likely that the interactions with different regulatory proteins, local chromatin environment or nearest neighboring factor binding motifs are likely important parameters underlying specific RA-transcriptional programs. Thus, identifying RAR and NR2F2 co-factors has the potential to shed light on the complex gene regulatory processes underlying CPC differentiation.

YAP1 and its homolog, TAZ, are the transcriptional effectors of the Hippo pathway; this pathway regulates cell proliferation and growth during development. When YAP1/TAZ are phosphorylated by Hippo kinases, they remain in the cytoplasm. When dephosphorylated, YAP1/TAZ translocate into the nucleus and interact with TEAD1-4 DNA binding proteins to induce expression of genes that promote proliferation^26^. While the importance of YAP1 in cardiomyocyte *proliferation* has been demonstrated^27,28^, only a few studies showed that YAP1 has a role in cell-fate decisions of the CPCs. For example, in zebrafish, knockout of the Hippo kinases increased the number of atrial cardiomyocytes, without affecting the proliferation of CPCs or the number of ventricular cardiomyocytes^29^. This suggests that YAP1 regulates atrial differentiation in the fish. However, the role of YAP1 in the differentiation of CPCs and the structure of the heart in mammals remains elusive.

In this study, we report that YAP1 is necessary for atrial lineage acquisition in human and mouse models of heart development. We found that RA signaling activates lineage enhancers containing YAP1 and TEAD4 proteins along with other key atrial specifiers, such as NR2F2. YAP1 deletion impaired RA-enhancer activation and altered the expression levels of lineage genes. Accordingly, we found that YAP1 is necessary for the patterning of the SHF and the expression of RA-target genes, such as *Nr2f2*, in gastrulating mouse embryos. Finally, YAP1 deletion severely compromises the formation of an atrial chamber in hESC-derived 3D cardiac patterned organoids, while the ventricle chamber is unperturbed. Overall, the data presented reveal that YAP1 is a novel factor integrated in the RA-dependent enhancer transcription factor network of CPCs necessary for the atrial differentiation route.

## RESULTS

### RA-induced transcriptome changes are captured in hCPCs

To gain insight into the mechanisms of RA in the specification of hCPCs, we utilized hESCs to model the developmental steps toward the acquisition of atrial and ventricular cardiac lineages. hESCs are differentiated to ventricular CMs following a temporal modulation of Wnt signaling (GiWi protocol^30^). However, the addition of 1uM RA to the media of early hCPCs, from Day 4 to Day 7, is sufficient to rewire the differentiation of hCPCs toward an atrial-like CM fate (**Figure 1A**). We performed single-cell transcriptome analysis at differentiation day 5 in the presence (RA^plus^) or absence of RA (RA^minus^) for 24h. We identified two predominant populations in our cultures, one corresponding to hCPCs (*PDGFRA*^+^)^31,32^ and the other one expressing high levels of endoderm markers such as *FOXA2* (*PDGFRA*^-^), which does not hold cardiac potential^32^ (**Figure 1B**). We found 159 differentially expressed genes (DEG) in RA^plus^ versus RA^minus^ hCPCs (64 upregulated and 95 downregulated, abs(Log2FC)≥0.5 and adj p<0.01) (**Table S1**). Untreated hCPCs expressed genes typically enriched in ventricular precursor genes such as ISL1^17^, whereas RA-treated hCPCs showed downregulation of these markers and increased levels of posterior-like SHF genes such as *MEIS2* and *HOXB1* genes^10,32^ (**Figure 1B**). Hence, our hESC-derived untreated and RA-treated hCPCs, express markers of ventricular and atrial progenitors, respectively. This is consistent with the developmental potential of these cells; as it is shown in **Figure 1C-D**, the RA^plus^ hCPCs preferentially differentiate into atrial-like CMs (*TNNT2*^+^ *MYH6*^high^ *NPPA*^high^ *NR2F2*^high^) compared to the RA^minus^ hCPCs, which mostly differentiate into ventricular-like CMs (*TNNT2*^+^ *MYL2*^high^ *LBH*^high^ *MYH7*^high^) (**Table S1**). We conclude that our cardiac induction strategy in hESCs effectively generated ventricular and atrial-like hCPCs and CMs in the presence and absence of RA, respectively.

**Figure 1.**
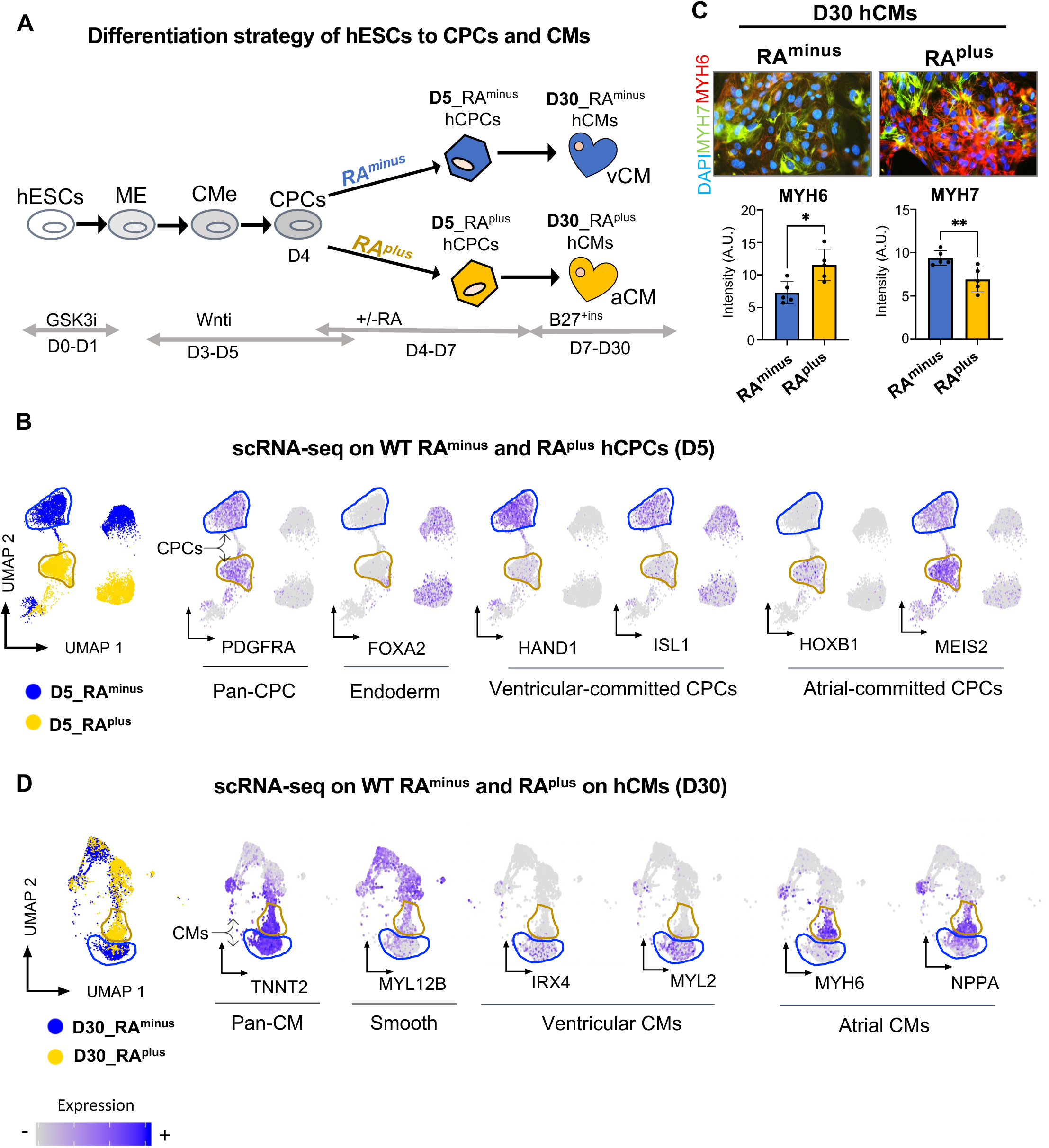
scRNAseq analysis of RA^minus^ and RA^plus^ hCPCs and hCMs captures the expression of atrial and ventricular lineage genes. **A)** Scheme depicting the strategy for cardiac induction in hESCs. 1uM of RA treatment (from day 4 to day 7) was used to induce atrial lineages. This study focuses on analysis of D5 CPCs and D30 CMs, untreated (RA^minus^) or treated with RA (RA^plus^). **B)** scRNAseq was performed on differentiation day 5 (D5) in RA^minus^ and RA^plus^ samples (1uM RA, 24h). UMAP plot showing indicated markers. Cells are colored by cell annotation (yellow and blue) or by expression of marker genes indicated below the graphs. **C)** Immunostaining and quantification of the atrial and ventricular markers MYH7 and MYH6, in D30 CMs, treated as indicated (n=5, Students t-test, *p<0.05, **p<0.01). **D)** scRNAseq was performed on differentiation day 30 (D30) in RA^minus^ and RA^plus^ samples (1uM RA, 72h from D4-D7). UMAP plots show markers of pan-CM, smooth muscle, ventricular, and atrial CMs. Two independent differentiation experiments were pooled for D5 and D30 sequencing experiments. Abbreviations: hESCs: human embryonic stem cells, ME: mesoendodermal, Cme: cardiac mesoderm, CPC: cardiac progenitor cells, RA: Retinoic Acid, v: ventricular, a: atrial, CM:cardiomyocyte, GSK3i: GSK3 inhibitor, Wnti: Wnt inhibitor, UMAP: Uniform manifold approximation and projection.

### RA signaling induces the opening of TEAD4-containing enhancers

To uncover novel mechanisms involved in the specification of hCPCs along with the RA signaling, we applied single-nuclei ATAC-sequencing technology^33,34^ to our RA^plus^ and RA^minus^ day 5 differentiation cultures. Similar to the scRNA-seq, two predominant hCPCs and endoderm populations were identified by differential chromatin accessibility of key lineage genes, including *TNNT2* (CPCs) and *FOXA2* (Endoderm) (**Figure S1A**). The RA treatment profoundly remodeled the chromatin accessibility profile of hCPCs: we found 11800 differentially activated regions (DARs) in RA^plus^ versus RA^minus^ hCPCs, while 7456 lost DNA accessibility (**Figure 2A**). Of these, 5586 and 4793 were associated to protein coding genes, respectively (**Figure 2A** and **Figure S1B**). Most of these genes were grouped in GO categories related to heart formation (**Figure S1C**), and a fraction of them showed changes in gene expression (**Figure 2B**).

**Figure 2.**
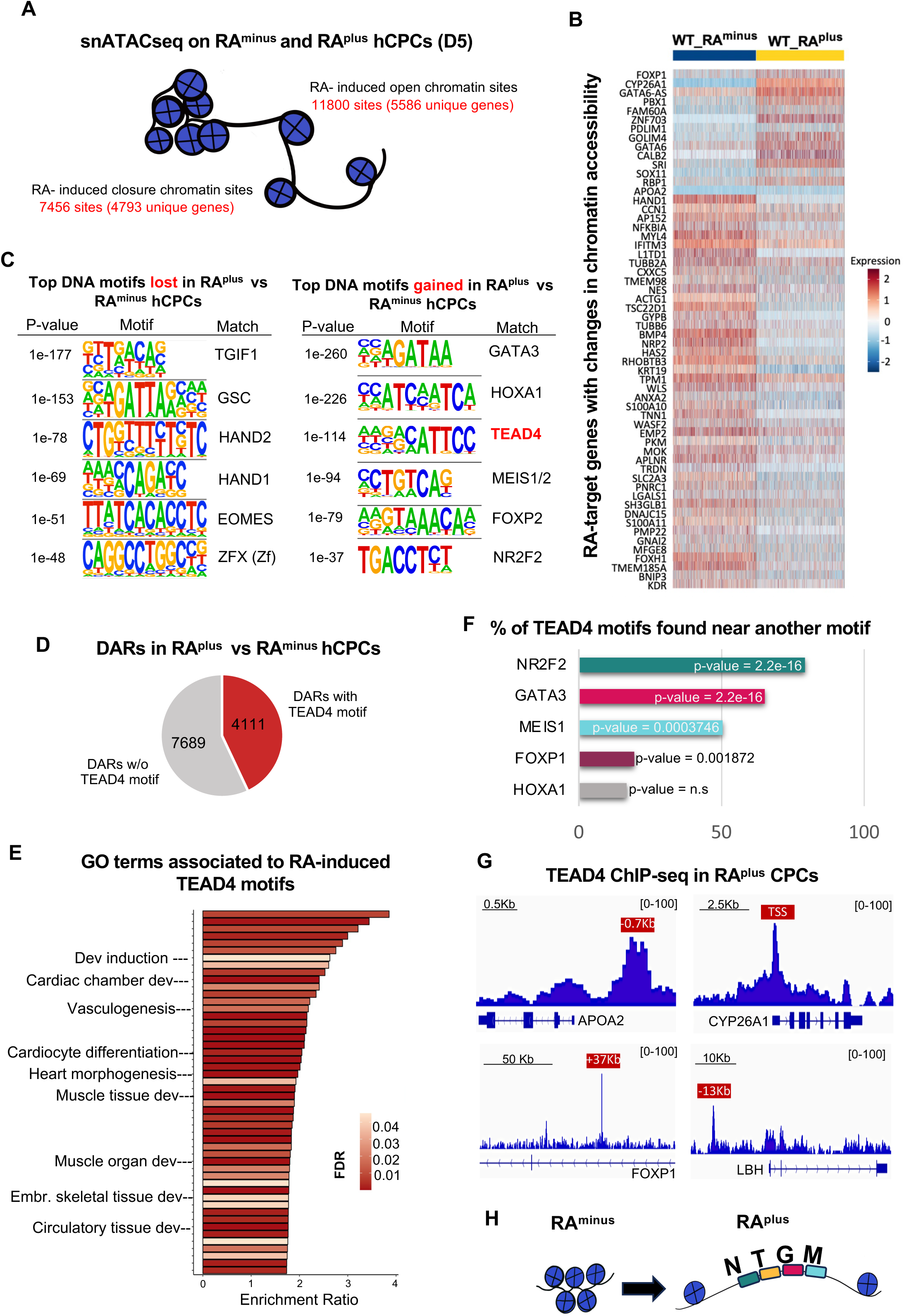
RA treatment in hCPCs triggers genome-wide opening of TEAD4 enhancers. **A)** snATACseq experiments were carried out in RA^minus^ and RA^plus^day 5 cultures (hCPCs). Scheme shows chromatin accessibility changes and associated number of genes induced by RA treatment in hCPCs. **B)** Heatmap of RA-target genes with correlative changes in chromatin accessibility (snATACseq) and gene expression (snRNAseq) in the hCPCs. **C)** De novo motifs analysis showing the top motifs lost and gained in RA^plus^ versus RA^minus^ hCPCs. **D)** Venn diagram showing the number of total RA-induced differentially activated regions (DARs) with and without TEAD4 motifs. **E)** Gene Ontology analysis of genes near RA-induced TEAD4 enhancers. **F)** Bars graph shows top DNA motifs found in the TEAD4 enhancers. The p value (Chi-square test for independence) indicates the motifs significantly associated to TEAD4. **G)** TEAD4 ChIP-seq was performed in RA-treated hCPCs. Genome browser captures show TEAD4 binding on RA-induced enhancers containing TEAD4 binding motifs. Red boxes show distance of RA-TEAD4 motifs from nearest TSS. **H)** Summarizing scheme showing that RA signal induces the opening of enhancers containing TEAD4 (T), NR2F2 (N), GATA3 (G) and MEIS1/2 (M) binding sites. Two independent differentiation experiments were pooled for snATACseq processing. Two independent ChIP experiments were pooled for sequencing.

We ran a DNA motif analysis through RA-regulated regions. Among the less accessible regions, TGIF1 binding motifs were the most impacted in response to RA (**Figure 2C**). TGIF1 is an inhibitor of the RA-dependent RXR alpha transcription^35,36^ and a transcriptional co-repressor of the TGFb-effectors SMADS^37^, and yet, its activities are important for specification and proper outflow tract septation^38^. The HAND1 and HAND2-recognized motifs also lost accessibility in RA-treated hCPCs, correlating with the downregulation of HAND1 mRNA levels observed in RA-treated cells (**Figure 2C** and **Figure1B**). Among the regions that gained accessibility, we found DNA motifs bound by TFs regulated by RA signaling in hCPCs, including MEIS1/2, HOXA1, and NR2F2^39^. Interestingly, we found that 4111 of the RA-opened sites contain the motif recognized by TEAD4 (the 3^rd^ motif most enriched, **Figure 2C**, **Figure 2D and Table S2**). Gene ontology (GO) analysis revealed that these RA-induced TEAD4 enhancers lie near genes controlling cardiac chamber development and cardiomyocyte differentiation (**Figure 2E**). Furthermore, 79.6% of RA-induced TEAD4 enhancers contain the motif recognized by the atrial TF NR2F2, suggesting that these factors form a transcriptional complex in response to RA. Other motifs significantly associated to TEAD4-motif containing enhancers were GATA3, MEIS1/2 and FOXP1 (**Figure 2F** and **Figure S1D**). To confirm the presence of TEAD4 protein in these enhancers, we performed ChIP-seq analysis of TEAD4 in RA-treated hCPCs. We identified 4679 TEAD4 peaks in the chromatin of RA-treated hCPCs and a significant association with RA enhancers containing TEAD4 binding sites (**Figure 2G** and **Figure S1E**). Altogether, these data show that RA induces lineage enhancers containing TEAD4 along with critical atrial transcription factors (**Figure 2H**).

### YAP1 binds and regulates RA-lineage enhancers containing TEAD4

In general, TEAD TFs associate to co-repressors in a close-chromatin conformation status and the binding of YAP1 is necessary to release the co-repressors and open the chromatin to regulate transcription^40^. We observed that RA triggers the opening of TEAD4-containing enhancers, thus, we speculate that RA induces YAP1 binding to these sites. To test this, we examined the binding of YAP1, TEAD4, and NR2F2, in the absence and presence of RA in the hCPCs by ChIP-qPCR analysis. **Figure 3A** shows that RA treatment triggers the recruitment of YAP1, along with NR2F2, to TEAD4 containing enhancers near *APOA2*, *APOA1* and *LBH* genes. Interestingly, RA treatment did not significantly affect binding of YAP1 to the canonical Hippo-target gene CTGF (**Figure 3A**). This suggests that RA specifically induces the binding of YAP1 to lineage enhancers. To examine the genome-wide occupancy of YAP1 to RA-activated enhancers, we performed ChIP-seq analysis of endogenous YAP1 protein in RA-treated hCPCs. Of a total of 6428 YAP1 peaks, 670 peaks associated to RA-induced regions (**Figure 3B**), and the majority of these regions contained TEAD4 sites (531/670, Chi-square test for independence p value:2.26^-16^) (**Figure S2A and Figure S2B**).

**Figure 3.**
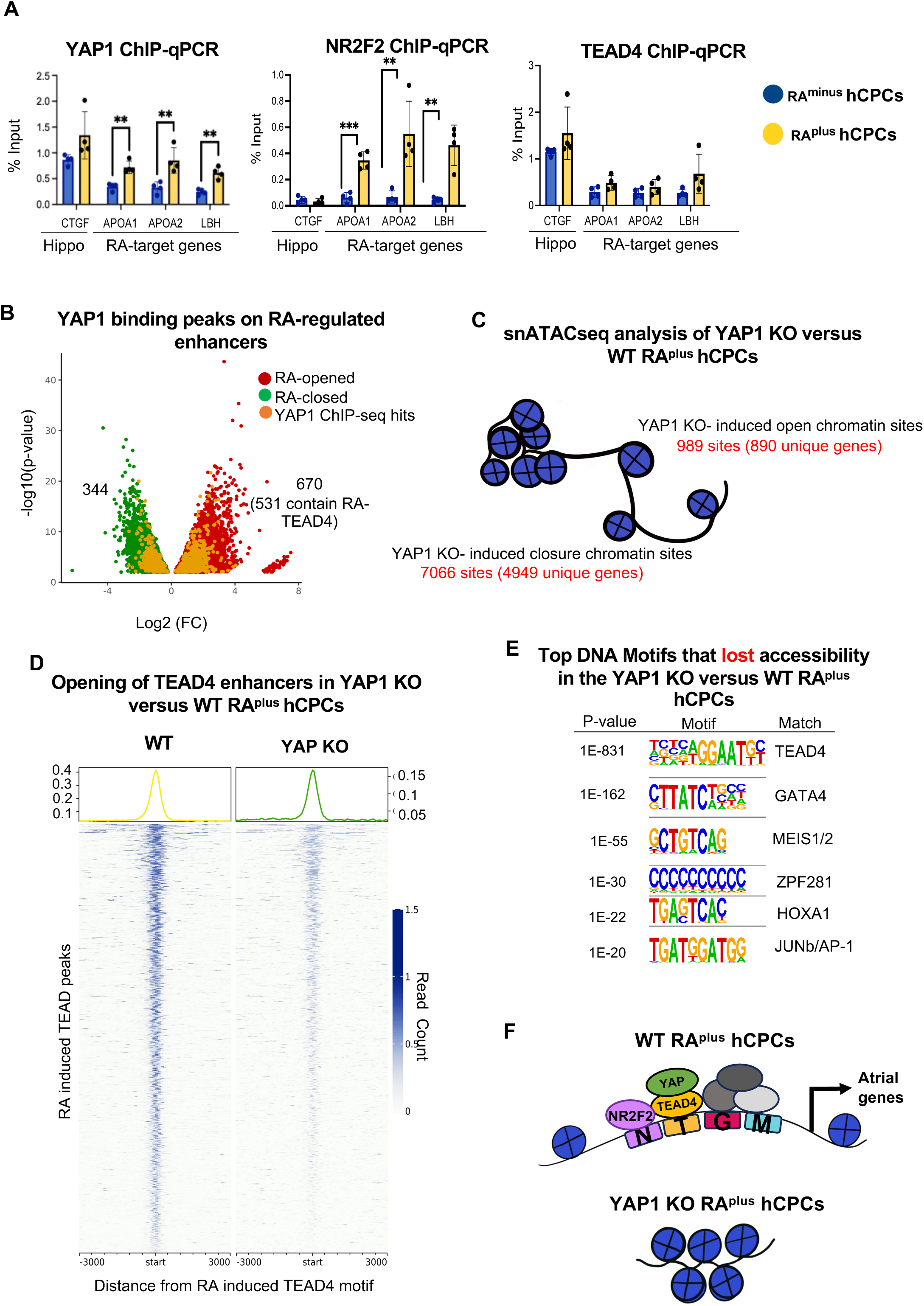
RA induces YAP1 binding to TEAD4 enhancers in hCPCs. **A)** ChIP-qPCR analysis of indicated proteins in RA^minus^ and RA^plus^ hCPCs. The analyzed gene regions are depicted in the X axis. Note that RA induces binding of YAP1 to TEAD4-enhancers on APOA1, APOA2 and LBH genes but does not affect binding to the Hippo target gene CTGF (n=4, p<**0.01, ***0.001). **B)** Volcano plot shows YAP1 bound peaks (from ChIP-seq; number of overlapping peaks are indicated) on RA regulated enhancers (snATAC-seq). **C)** snATACseq was performed in RA-treated YAP1 KO hCPCs and compared to WT. Scheme depicts the number of chromatin regions that change accessibility in the absence of YAP1. **D)** Heatmap shows the chromatin accessibility (peak intensity) of RA-induced TEAD4 enhancers in WT and YAP1 KO RA^plus^ hCPCs. **E)** Top motifs found in the DNA motif analysis of the 7066 regions that lost accessibility in the RA-treated YAP1 KO hCPCs compared to WT. **F)** Summarizing scheme shows that YAP1 is necessary for the opening of RA-enhancers in hESC-CPCs. Two independent differentiation replicates were pooled for sequencing.

To interrogate whether the opening of RA-induced TEAD4 sites in hCPCs are YAP1-dependent, we performed single-cell ATACseq experiments in RA-treated YAP1 KO hCPCs (Day 5), and compared to WT. Globally, we identified that 7066 DNA regions lost accessibility in YAP1 KO compared to WT hCPCs, while 989 regions gained accessibility, consistent with a general role of YAP1 in facilitating chromatin opening (**Figure 3C**). Among the RA-induced chromatin regions, 1722 lost access in the YAP1 KO, from which 1275 (75%) contained TEAD4-motifs activated regions (**Figure 3D and Figure 3E**). Furthermore, the GO analysis of the RA and YAP1 regulated chromatin regions showed enrichment on critical cardiac development terms, including cardiomyocyte differentiation (**Figure S2C**). Overall, these findings show that YAP1 is necessary for the opening of critical RA-enhancers in hCPCs (**Figure 3F**).

### YAP1 regulates RA-target genes in hCPCs

We analyzed how YAP1 levels affect the transcriptome of the RA-treated hCPCs by scRNAseq analysis. Similar to WT, we identified two main populations corresponding to CPCs and endodermal cells in the YAP KO D5 cultures (**Figure 4A**). Our analysis showed that YAP1 deletion affected the expression levels of 74 genes (41 upregulated and 33 downregulated, abs(Log2FC)≥0.5 and adj p<0.01, and **Figure 4B** and **Table S3**). Among these, we identified the typical *NR2F2*-target genes *APOA1* and *APOA2*^41^, which are also direct YAP1-targets (**Figure 3A** and **Figure S2A**), and other RA-target genes involved in heart development, including *KDR*^42^, *CHCHD10*^25^, *GYPB*^43^ and *HAS2*^44^ (**Figure 4B**). We also found specific atrial genes downregulated in the YAP1 KO hCPCs, including *MYL7*^45^, *ACTC1* and *NPPB*^46–48^. On the other hand, ventricular lineage genes, such as *HAND1*^49^ and *MEF2C*^50^ were upregulated in the YAP1 KO hCPCs (**Figure 4B** and **Figure 4C**), suggesting that YAP1 is needed to silence the ventricular program. Furthermore, unbiased GO analysis of DEGs revealed multiple categories related to heart formation, further supporting the relevance of YAP1-target genes for CPC differentiation (**Figure 4D**). In addition, our ChIP-seq analysis revealed YAP1 binding peaks near *MYL7* and *NPPB*, suggesting that YAP1 directly activates these atrial genes, in agreement with its general association to RA-induced enhancers. However, we do not detect YAP1-binding peaks on *HAND1* or *MEF2C* genes, suggesting that these YAP1-repressed genes are indirect targets (**Figure 4E)**. Altogether, these results show that YAP1 is necessary to execute the RA-transcriptional program in hCPCs to promote the acquisition of atrial lineages and turn down alternative cell fates (**Figure 4F**).

**Figure 4.**
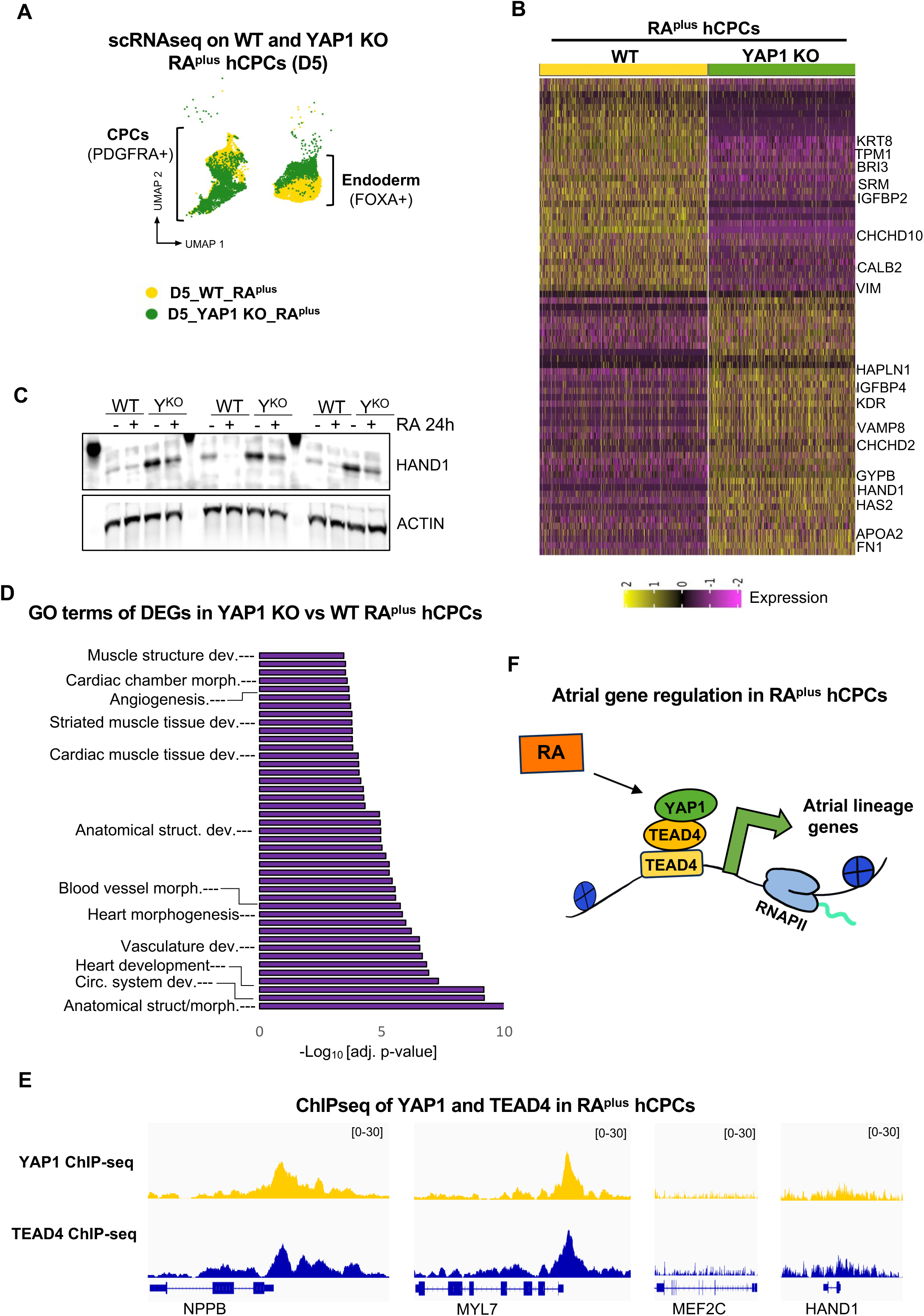
YAP1 regulates lineage genes in RA^plus^ hCPCs. **A)** UMAP distribution of RA^plus^ WT and YAP1 KO Day 5 cultures, colored by sample identity. The two main populations identified corresponding to endoderm (Foxa2+) and CPCs (Pdgfra+) are highlighted. **B)** Heatmap shows the differentially expressed genes in the YAP1 KO CPCs versus WT from scRNAseq analysis (Log2 FC>[0.5], adj p<0.01). **C)** Western Blot shows expression of HAND1 in WT and YAP1 KO D5 CPCs untreated (-) or treated with RA for 24h (+). Actin was used as loading control. Three biological independent replicates are shown. **D)** GO analysis of DEGs in the YAP1 KO vs WT RA^plus^ hCPCs. Heart development-related categories are highlighted. **E)** YAP1 ChIP-seq analysis were performed in WT RA^plus^ hCPCs. Genome browser captures show YAP1 binding distribution on the indicated genes. Two independent ChIP experiments were pooled for sequencing.

### YAP1 compromises the identity of atrial-like CMs in directed differentiation approaches

Our findings above suggest that YAP1 regulates the differentiation of atrial CMs. To gain insight into this idea, we differentiated RA-treated YAP1 KO hCPCs toward atrial like CMs (aCMs), using 2D differentiation approaches (**Figure 1A**). RA-treated YAP1 KO hCPCs differentiated into beating CMs to a similar rate as controls (**Figure S3A**). However, scRNAseq analysis revealed critical developmental genes differentially expressed in the YAP1 KO aCMs compared to WT (**Figure S3B, Figure S3C,** and **Table S3)**. These genes include *HES1*, a NOTCH target gene expressed in the SHF, important for development of the outflow tract^51^, and *ACTA1*, which encodes for skeletal muscle alpha-actin and has been implicated in myopathies affecting the heart^52^. Following the same trend than the hCPCs, YAP1 KO aCMs expressed higher levels of genes typically enriched in ventricular CMs, such as *LBH*^53^, compared to WT. Convergently, the expression of common atrial markers, such as *NPPA* and *KCNA5*^46,47^ were reduced in the YAP1-deleted aCMs (**Figure S3B**). We also identified a group of genes involved in Ca^+2^ signaling and mitochondria, including Phospholamban (PLN), the prototypical negative regulator of SERCA^54^, and *CHCHD2*, a regulator of oxidative phosphorylation and mitochondrial stress^55^ (**Figure S3B and Figure S3D**). To further assess the phenotypic properties of the WT and YAP1 KO atrial-like CMs, we performed calcium imaging using the radiometric reporter, Fura4. Interestingly, we detected a significant difference in the peak amplitude of the Ca^+2^ transients in the YAP1 KO atrial-like CMs when compared to WT (**Figure S3E**). Altogether, these results indicate that YAP1 is necessary for the activation of the atrial program and the acquisition of an atrial CM identity.

### YAP1 is necessary for the development of an atrial chamber in 3D patterned cardiac organoids

To further dissect the role of YAP1 in heart chamber development, we incorporated a pioneer model of hESC-derived cardiac organoids^16^. These organoids develop from hESCs in a self-assembly fashion and develop distinct atrial and ventricular chambers, facilitated by the activity of an endogenous RA gradient^16^. To investigate the role of YAP1 in the patterning of these structures, we generated cardiac organoids derived from control and YAP1 KO H1 hESCs (**Figure 5A**). YAP1 KO organoids develop effectively from Day 0 to Day 30, but morphological analysis reveal that the KO organoids are more elongated than WT (**Figure 5B**, **Figure 5C**, and **Figure S4A**). We analyzed the formation of the atria and ventricle by examining the expression domains of well-defined markers restricted to atrial and ventricular chambers, *MYL3*^16^ and *NR2F2*^16,18,19^, respectively. Analysis of day 30 organoids revealed that the size of the ventricle chamber is similar in WT and YAP1 KO organoids, as indicated by the area marked by *MYL3*. Strikingly, the area of the atrial chamber, measured by the expression domain of *NR2F2*, is significantly reduced in the YAP1-deleted organoids (**Figure 5D** and **Figure 5E**). We conclude that YAP1 is necessary for the formation of an atrial chamber in the hESC-derived cardiac organoids.

**Figure 5.**
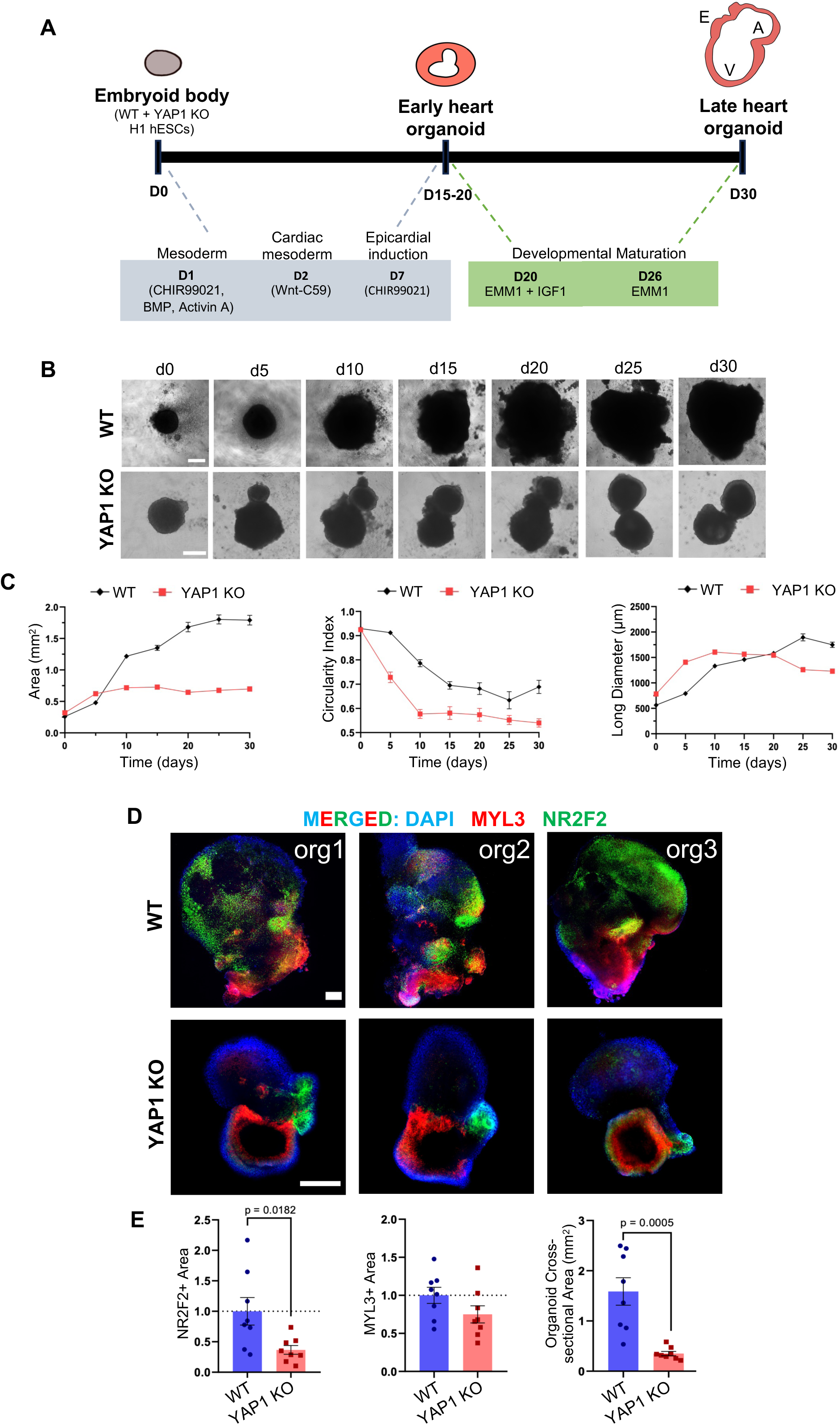
YAP1 is required for atrial chamber development in hESC-derived patterned cardiac organoids. **A)** Scheme outlines the cardiac induction protocol, adapted from Volmert at al., Nat Comm. 2023. The D30 organoids have (A) atrial, (V) ventricular, and (E) proepicardium structures, as indicated in the scheme. **B)** WT and YAP1 KO H1 hESCs were differentiated to cardiac organoids as shown in (A). Representative bright field images are shown, taken at the indicated days. **C)** Graphs show quantification of the area (left), shape (middle) and length (right) of the WT and YAP1 KO organoids on the indicated days of differentiation. Two differentiation experiments were done with similar results and 7-8 organoids were analyzed. **D)** Immunofluorescence images of three representative WT and YAP1 KO day 30 organoids stained with DAPI (blue), the ventricular marker MYL3 (red), and the atrial marker NR2F2 (green). Scale bars, 200µm **E)** Quantification of NR2F2+ and MYL3+ areas and organoid cross-section area. Students t-test (n=7-8, Students t-test).

### Efficient depletion of YAP1 levels in embryonic clusters of late gastrulating embryos

Recent evidence suggest that the distinct cardiac progenitor populations are specified as they emerge from the differentiating epiblast cells of the gastrula^56,57^. Thus, to interrogate the role of YAP1 in cardiac specification in vivo, we used a *Yap1^Flox/Flox^:Sox^cre^* system to conditionally delete *Yap1* in the epiblast^58^ (**Figure 6A**). *Sox2* is expressed in the pluripotent cells that undergo gastrulation, generating the three germinal layers; ectoderm, endoderm, and mesoderm. Among these, the mesoderm layer harbors the different hCPCs pools, that migrate toward the anterior part of the embryo as they emerge from the Primitive Streak (**Figure 6A**). We performed scRNAseq analysis on E7.75 embryos, which by our eye staging corresponds to a late streak (**Figure S5A**).

**Figure 6.**
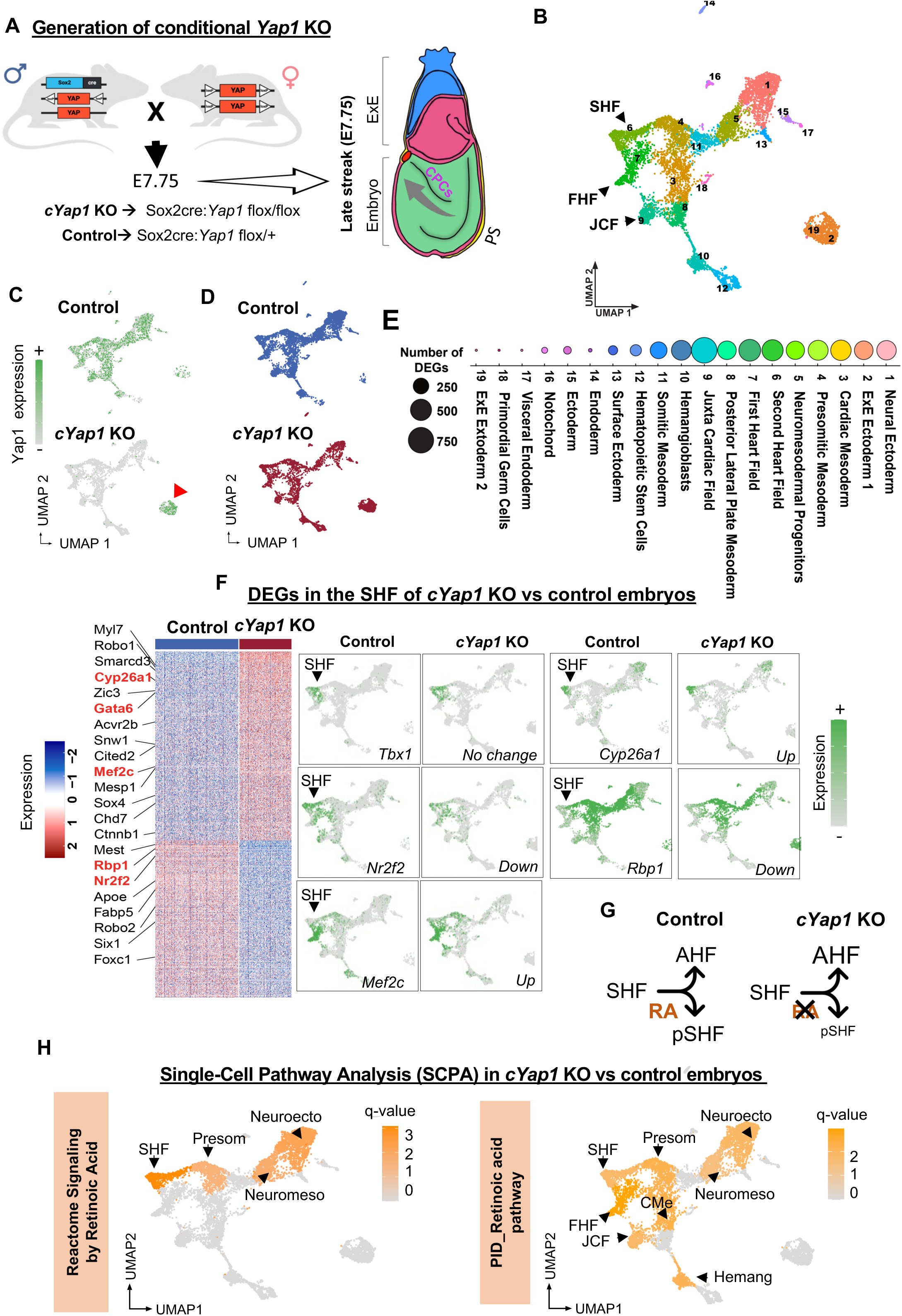
*cYap1* deletion in vivo alters the transcriptome of the SHF and Vitamin A/RA pathway. **A)** Breeding scheme used to generate *cYap1* KO and heterozygous control embryos, and a simplified scheme of a E7.75 embryo. **B)** scRNAseq was performed on 10 *cYap1* KO and 7 control E7.75 embryos. Unbiased clustering identified 19 clusters in these embryos, that were annotated as indicated in **6E** using known markers (see also **Figure S5E)**. **C)** UMAP showing Yap1 expression levels in control and cKO embryos. Red arrowhead marks the extraembryonic (ExE) (Sox2^-^, Rhox5^+^). **D)** UMAP shows space distribution of normalized number of cells in control and cYap1 KO embryos. **E)** Dotplot graph showing the number of DEGs in *cYap1* KO vs control embryos in each cluster. Note that the bigger the dot the more DEGs in a given population (abs(Log2FC)>0.25, adj. p value<0.05). **F)** Heatmap shows DEGs in the SHF of *cYap1* KO vs control embryos. Genes relevant for heart development are highlighted. Genes in red are directly involved in SHF patterning and RA activity. UMAPs showing the expression levels of critical SHF genes. **G)** Scheme summarizing the effect of Yap1 depletion in the SHF. **H)** UMAPs showing results from unbiased SCPA analysis on Vitamin A/RA signaling terms indicated in the vertical orange boxes. Significant changes in pathway activity are displayed in orange color. The grey color indicates no enrichment. The name of the populations with significant changes are indicated.

Following optimized procedures in our lab^59^, 10 *cYap1* KO (*Yap1^Flox/Flox^:Sox^cre^*) and 7 control (*Yap1^Flox/+^:Sox^cre^*) embryos were processed for sequencing (**Figure S5B**). Bioinformatics analysis confirmed high-quality sequencing results (∼4K cells/sample and ∼30K reads/cell and **Figure S5C and S5D**). We annotated 19 clusters based on the expression of known markers, including the three main CPC populations: FHF (*Tbx5^+^*), SHF (*Tbx1^+^*) and the recently discovered Juxta Cardiac Field (JCF, *Mab21l2^+^*)^60^ (**Figure 6B, Figure S5E, and Figure S6A**). Importantly, *Yap1* deletion was confirmed in all *Sox2*^+^-derived clusters in the c*Yap1* KO embryos (**Figure 6C**).

### Conditional YAP1 deletion does not affect the developmental timing of cardiac trajectories

We then projected *cYap1 KO* cells onto these control cell populations. All *cYap1 KO* cells could be projected to control cell populations, with each control population receiving some *cYap1 KO* cells (**Figure 6D** and **Figure S6B**), but in varying proportions. The pseudotime analysis of E7.75 embryos identified 8 trajectories originating from a common founder, the epiblast (that at E7.75 is committed to neuroectoderm, cluster 1) (**Figure S6C**). Trajectory 1, corresponding to the hematopoietic lineage, emerged from the first primitive streak population, and showed accelerated differentiation in *cYap1* KO embryos, correlating with an increase in hematopoietic precursors (cluster 12) in the *cYap1* KO compared to controls (**Figure S6B** and **Figure S6C**). Trajectories 2 to 4, representing other primitive streak-derivatives, including cardiac and other mesoderm lineages, showed slight acceleration in differentiation, although to a lesser extent than trajectory 1. Accordingly, the number of cells in the three main heart fields remained similar between control and *cYap1* KO embryos, which facilitates precise comparison of gene expression levels (**Figure S6B**). We observed a reduction in the number of *cYap1* KO cells in the epiblast and surface ectoderm, corresponding to trajectory 8, that exhibited delayed differentiation in *cYap1* KO embryos (**Figure S6B and S6C**). We conclude that deletion of *Yap1* in the *Sox2^cre^*lineage accelerates the differentiation of epiblast cells toward primitive streak-derived lineages, and earlier differentiation trajectories (i.e hematopoietic) are the most differentially affected by the number of cells.

### Yap1 regulates the transcriptome of the cardiac fields and the patterning genes of the SHF

During cardiogenesis, three main heart fields contribute to the formation of the heart; the FHF, SHF and JCF. The SHF is the main contributor of atrial cells, and it is patterned into anterior and posterior populations before differentiating into final cells. The anterior SHF (AHF) progenitors are characterized by the expression of *Mef2C*, *Fgf8*, and *Fgf10* and contribute to right ventricular cardiomyocyte (RVCM) and outflow tract (OFT) lineages^14,61^, whereas the posterior SHF (pSHF) progenitors, defined by the expression of *Hoxa1, Hoxb1, Hotairm1* and *Nr2f2*, give rise to atrial CMs and sinus venosus-derived structures including the sinoatrial node pacemaker cells^10,19,62^. We investigated whether the decision to adopt an anterior or posterior lineage identity may be disrupted upon *Yap1* loss. Among the 19 clusters identified, the three heart fields were the ones with the highest number of differentially expressed genes in the *cYap1* KO , compared to control embryos (**Figure 6E and Table S4**). Among the 590 DEGs in the SHF ( abs (Log2FC)>0.25, adj. p val <0.05), 322 were upregulated and 268 downregulated (**Figure 6F**). Many of the DEGs, including *Six1*^63^, *Cited2*^64^, *Smarcd3*^65^ and *Chd7*^66^, are involved in the formation of heart structures derived from the SHF progenitors; such as the outflow tract, arteries, and septum (**Table S5**). Importantly, we also found DEGs directly involved in the patterning of the SHF cells. The pSHF markers *Nr2f2* and *Hotairm1* were downregulated, and the AHF markers *Gata6*^67^, *Mef2c*^50,68^ and *Irx5*^69^ were upregulated in the *Yap1* KO SHF cells (**Figure 6F**). Overall, we conclude that *cYap1* deletion in the SHF cells impairs the induction of posterior lineage genes, such as *Nr2f2*, which likely compromises the differentiation of the SHF and acquisition of atrial lineages during in vivo development (**Figure 6G**).

### Yap1 regulates the activity of the RA signaling pathway in the embryo

The activity of RA signaling is necessary for the acquisition of a posterior SHF lineage. Thus, we performed a single-cell pathway enrichment analysis (SCPA^70^) focused on the Vitamin A/RA pathway in *cYap1* KO versus control embryos. Interestingly, Vitamin A/RA pathway genes were significantly enriched across the clusters of the cardiovascular trajectories, including the cardiac mesoderm, the FHF and SHF (**Figure 6H**), along with the neuroectoderm and somitic mesoderm upon *cYap1* deletion. Accordingly, the RA-target gene Hotairm1 was downregulated across several of these clusters in the cYap1 KO embryos, while *Hdac1* and *Ncoa2*, typical RAR corepressors, were upregulated (**Figure S6D and Table S4**). We also observed DEGs involved in the overall activity of the Vit A/RA pathway, that regulate the levels of RA or its cellular distribution. For instance, in the SHF, *Rbp1*, which transports Retinol inside the cell^71^ was low in *cYap1* KO, compared to controls, and *Cyp26a1*, the enzyme responsible for RA degradation^72^ was upregulated (**Figure 6F**). We also observe a downregulation *trend* in *Aldh1a2* levels, the enzyme responsible for RA production, across clusters (**Figure S6D**). Altogether these results show that *Yap1* deletion alters Vit A/RA metabolic genes and suggests that it may decrease the bioavailability of RA in the early embryo. These conclusions are further supported by the increase in the levels of the RA-repressed gene, *Hand1*, in the nascent mesoderm of cYap1 KO embryos compared to controls (**Figure S6E**), following the same regulation that we observed in vitro (**Figure 4C**). Overall, we conclude that *Yap1* regulates both RA-target genes involved in lineage acquisition and pathway homeostasis.

## DISCUSSION

During heart development, RA signaling acts on the CPCs of the SHF to specify an atrial CM fate, a critical step in determining the identity of the heart chambers. In this study, we discovered that RA-induction of the atrial CM program requires the Hippo effector YAP1. Our findings demonstrate that RA treatment triggers *YAP1* recruitment to cardiac enhancers alongside with the atrial specifier *NR2F2*. The deletion of *YAP1* hinders the retinoic acid (RA)-induced opening of enhancers and expression of critical atrial genes. In agreement, we also found that *YAP1* is necessary for the formation of an atrial chamber in hESC-derived organoids, while it is dispensable for ventricle development. Our in vivo analysis further supports these findings by showing that *YAP1* regulates the activity of Vitamin A/RA in the early embryo and the anterior-posterior patterning of the SHF (**Figure 7**).

**Figure 7.**
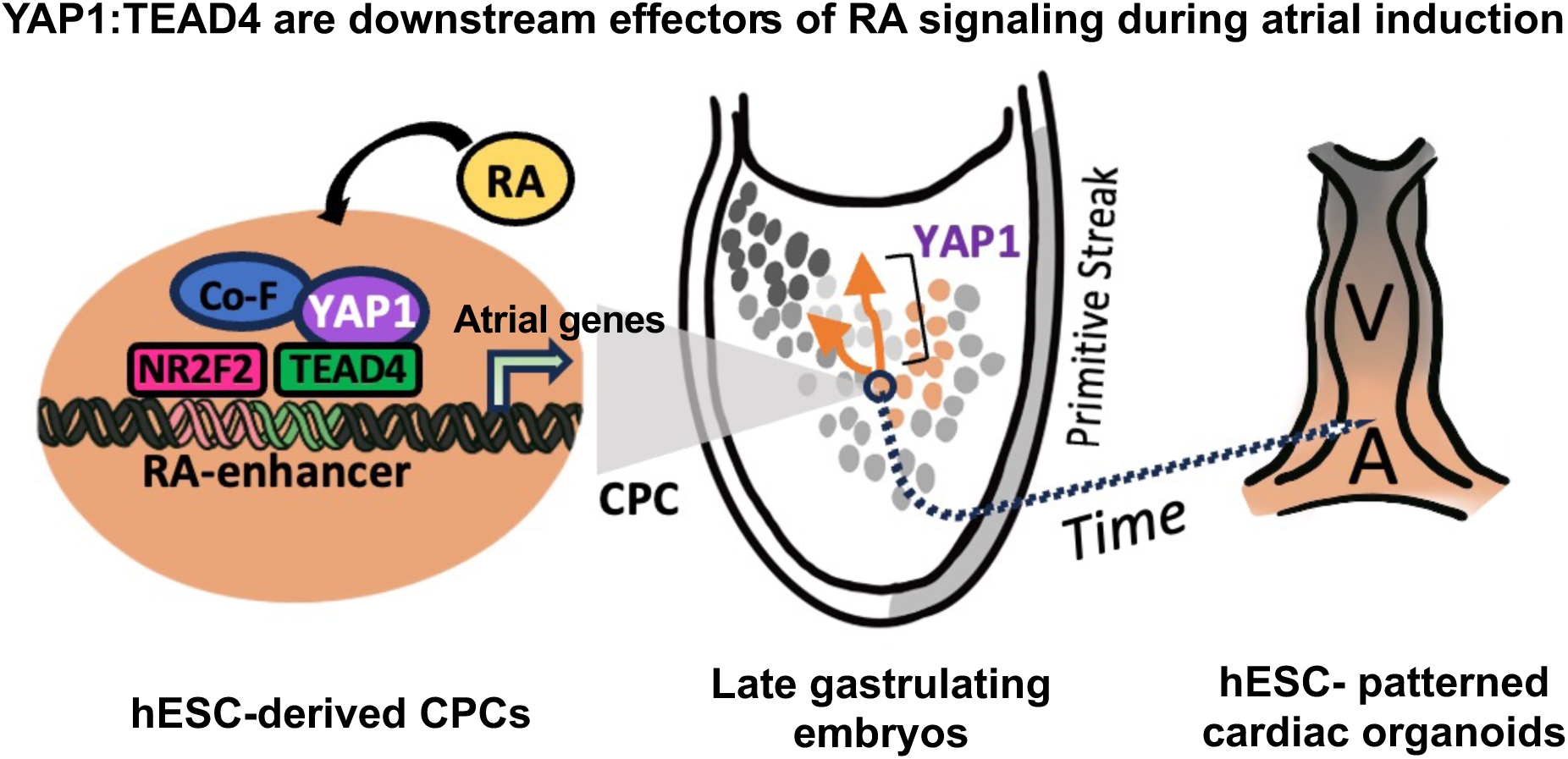
Scheme summarizing results. YAP1 is a downstream effector of Vit A/RA signaling essential for the acquisition of atrial lineages. RA signaling induces YAP1 binding to TEAD4 and NR2F2-containing enhancers in CPCs, and YAP1 opens the chromatin and activates atrial genes (on the left, hCPCs). The activity of YAP1 is necessary for the acquisition of posterior trajectories in CPCs (“orange dots”, embryo model in the middle) and the formation of their derived structures, such an atrial chamber (on the right, organoid models).

Our discovery that YAP1 is a non-canonical effector of the RA signaling pathway in CPCs started with the observation that RA treatment induced a massive opening of TEAD motifs across the chromatin of human ESC-derived CPCs. Our findings are consistent with previous observations in mice embryos, showing that enhancers containing TEAD4 motifs are enriched in the posterior second heart field (pSHF) (the *in vivo* counterpart of the RA-treated hCPCs), compared to anterior second heart field (AHF)^62^. However, the role of these enhancers had not yet been studied. Our studies build upon this observation to reveal that the RA signaling is the input that induces the opening of the TEAD motifs in the pSHF cells, through the induction of YAP1 recruitment to these sites. Moreover, we provide evidence, in vitro and in vivo, that the functional role of the YAP1:TEAD enhancers is to regulate critical RA-target genes important for the acquisition of atrial lineages.

Our findings show that up to 79% of RA-induced TEAD4 enhancers also contain NR2F2 motifs, and we confirmed the RA induces the binding of NR2F2 protein, along with YAP1, to a set of these enhancers. However, we did not resolve the transcription factor hierarchy of these enhancers. While TEAD4 TFs are already bound to the enhancers in the absence of stimuli (in this case RA), whether NR2F2 brings YAP1 or vice versa to open the enhancers is an unresolved question. A recent publication reported a physical interaction between NR2F2 and YAP1 proteins in prostate cancer cells^73^. Moreover, other nuclear receptors, including the estrogen receptor^74^ and the own RA-receptor RAR^75^, can recruit YAP1 to the chromatin in cancer cells. Thus, it is possible that NR2F2 binding to these enhancers unwinds the chromatin to expose TEAD proteins favoring subsequent YAP1 binding or directly recruits YAP1 through a physical interaction. These hypotheses are currently being tested in our lab.

NR2F2 determines the identity of an atrial chamber during heart development^19^. Given that YAP1 and TEAD4 were tightly associated to NR2F2-containing enhancers in RA-treated CPCs, we examined whether YAP1 is necessary for the formation of an atrial chamber. We tested this possibility in human cardiac organoids, a model that partially recapitulates the development of an atrial and ventricular chambers. We found that YAP1 is essential for the development of an atrial chamber (NR2F2+), while is dispensable for the ventricle (MYL3+) in the human organoids. These results highlight the relevance of YAP1 working as a RA-effector and regulator of atrial lineage genes. Given that the CPCs of the SHF give rise to other heart cells besides atrial CMs, including those of the outflow tract, more studies are needed to investigate whether YAP1 regulates the formation of other structures of the heart.

Our findings are pioneer revealing a fundamental role of YAP1 controlling cell-fate decisions during early heart development. Previous studies in mouse embryos reported that conditional YAP1 deletion in the cardiac lineage leads to hypoplastic hearts without significantly affecting overall heart structure, suggesting that YAP1 does not control differentiation of CPCs^27,76^. To the best of our knowledge, the earliest ablation of YAP1 in the cardiac lineage in mice was done using a knock-in *Nkx2.5*-cre mouse line, which drives *Yap1* deletion at E7.5 in the cardiac progenitors of the cardiac crescent^77^. This *cYap1* KO impaired cardiomyocyte proliferation, leading to severe myocardium hypoplasia and embryonic lethality^27^. However, under these genetic conditions, the structure of the heart was preserved, suggesting that *Yap1* deletion did not affect the differentiation of the cardiac progenitors. However, this model is not suitable for studying the role of YAP1 in CPCs for a few reasons. First, the *Nkx2.5*-cre driver becomes active at ∼E7.5, which is too late, as evidence show that CPCs commit to their lineages during gastrulation at ∼E6.5^56,78^. Second, the *Nkx2.5*-Cre driver is mostly active in CPCs committed to ventricle lineages^77,79^, as other CPCs (atrial committed CPCs^11,80^ or CPCs from the Juxta Cardiac Field^60^) do not express *Nkx2.5*. Thus, additional mouse models, including cre-drivers that delete *Yap1* earlier and, in all CPCs, were necessary to understand the role of YAP1 in CPC biology. Using a *Sox2*-cre driver, that deleted YAP1 around E5.5 in the pluripotent cells of the epiblast, we uncovered new YAP1-target genes in the CPCs. Among them, we identified *Nr2f2* and *Mef2c* genes, master regulators of SHF patterning and chamber development^19,68^, which strongly support the notion that YAP1 participates in the cell-fate decisions of the CPCs, in vivo.

We found that YAP1 regulates transcription of RA-target genes involved in (**I**) CPC commitment and (**II**) Vit A homeostasis. Vit A homeostasis refers to the capacity to maintain the pathway’s activity despite a range of fluctuations in substrate availability. Mutations in the genes that regulate RA production or activity disrupt Vit A homeostasis and impact heart development. These include family members of ALDH^81^ (RA producers) and CYP26 enzymes^72,82^ (RA degraders), CRBPs^71^ (RA transporters) and downstream effectors, like NR2F2^22,23^. Furthermore, Vit A activity can also be impacted by mutations in cardiac genes. A vivid example is the TBX1 gene, whose haploinsufficiency causes heart phenotypes in DiGeorge Syndrome^83,84^. Mice lacking *Tbx1* have increased expression of ALDH1A2 and reduced expression of CYP26 enzymes, leading to an accumulation of RA, and phenotypes resembling excess of Vit A^85,86^. Our data revealed that *Yap1* depletion significantly alters the expression of key regulatory Vit A/RA enzymes, including *Cyp26a1*. These observations lead us to speculate that genetic conditions, such as TBX1 haploinsufficiency, may disrupt Vit A homeostasis by influencing YAP1 activity. Thus, it would be interesting to investigate the contribution of YAP1 to Vit A dysregulation in patients with CHD, such as DiGeorge syndrome.

Overall, our findings define a new signaling axis composed by the Vitamin A/RA and Hippo/YAP1 signaling pathways, that is essential for the acquisition of atrial lineages and formation of an atrial chamber. Our findings have implications for heart development and CHD. Furthermore, the revelation of YAP1’s role in the patterning of the atrial and ventricular chambers could be leverage for advancing organoid engineering by using small molecules to modulate YAP1 activity^87^ to precisely steer chamber development.

## Methods

### hESC culture and directed cardiac differentiation

All experiments were performed using WT and YAP1 KO H1 hESCs previously described^88^. These cells were cultured in mTESR1 media on 6-well Matrigel-coasted tissue culture plates. hESC colonies were split with 0.5mM PBS/EDTA to maintain stocks of the cells. Typically, cells were spilt at a 1:20 ratio every 5-7 days. For experiments, colonies were dissociated using Accutase and Rock inhibitor and counted with an automated cell counter per desired cell number. The differentiation protocol was adapted from previous publications^30,89^. Briefly, hESCs were passaged using trypsin or Accutase, and started differentiation when reached ∼80% confluency. At day 0, hESCs were treated with 6 µM ChIR in double feed of mTESR1 media. The timing of the day was recorded so that media could be replaced exactly 24 hours later. At day 1, exactly 24 hours later, the media were replaced with RPMI/B-27 minus insulin. On day 3, half the media was removed from each well from plate, mixed with an equal amount of fresh RPMI/B27 minus, and 7.5µM IWP-2 was added. RA 1uM was added on day 4 for atrial induction. On day 5, media was changed to RPMI/B27 minus insulin. The CPC analyses were done on day 5 cultures. From day 7 onwards, media were exchanged for RPMI/B27 plus insulin and replenished every 3 days. Contraction of cells begin from day 8 to 10. The analysis of atrial and ventricular CMs were performed at day 30 of differentiation.

### Immunostaining staining and 2D imaging of hESCs-CMs

hESCs-derived CMs were dissociated using Accutase and plated in chambered glass slides coated with laminin. Cardiomyocytes were recovered in culture for 3-5 days before continuing. Cells were fixed with 4% formaldehyde for 15 minutes at room temperature and wash two three times with DPBS-/-. After fixation, cells were permeabilization with 0.5% Triton in DPBS-/-for 20 minutes at room temperature. Cells were washed with DPBS-/-three times. Then incubated with blocking solution (3% Donkey serum in PBST) for 30 minutes at room temperature. Primary antibodies were diluted in blocking buffer (see **Table S6**) and incubated overnight at 4°C. The next day, the cells were washed 3 times with DPBS-/- and incubated with 1µg/ml DAPI for 10 minutes. Images were captured by a Nikon Eclipse Ti confocal microscope using Nikon NIS-Element AR software.

### Western Blotting

For whole-cell protein analysis, samples from untreated and RA-treated CPCs were lysed with RIPA buffer supplemented with protease and phosphatase inhibitors. The samples were then placed on ice for 10 minutes and centrifuged at 5000g for 10 minutes. The supernatant containing the protein was collected. The protein concentration was quantified using the Bicinchoninic acid (BCA) assay. Subsequently, protein samples (25ug) were loaded onto polyacrylamide Tris-glycine SDS gels and subjected to electrophoresis. Wet transfer was performed, and the gels were transferred to 0.45um PVDF membranes. These membranes were blocked for 1 hour at room temperature in 2% BSA in TBST, followed by overnight incubation with the primary antibody at 4 °C. The next day, membranes were washed three times for 5 minutes with TBST and then incubated with secondary antibody for 1 hour at room temperature. After another three washes for 5 minutes each with TBST, chemiluminescence detection was used, and the membranes were imaged on a Chemidoc system. The antibodies utilized in the study are listed in **Table S6**.

### Generation of hESC-derived patterned cardiac organoids

H1 hESCs (WT and YAP1 KO) were cultured in Essential 8 Flex Medium with 1% penicillin streptomycin (Gibco) in 6-well plates on growth factor reduced Matrigel (Corning) inside an incubator at 37 °C and 5% CO_2_. hESCs were passaged using ReLeSR passaging reagent (STEMCELL) upon reaching 60-80% confluency. Organoid formation comprised two phases. From Day 0 to Day 20, cardiac bodies were generated as previously described^90^. In the second phase, cardiac spheres were induced to maturation using the EMM2/1 developmental induction media from Day 20 to Day 30 as shown previously^16^. Cardiac organoids were processed at Day 30 for downstream analysis.

### Immunofluorescence, confocal microscopy, and image analysis of organoids

hESC-cardiac organoids were transferred to 1.5 mL microcentrifuge tubes using a cut 200 μL pipette. Organoids were fixed in 4% paraformaldehyde in PBS for 30 minutes. Following this, organoids were washed using PBS-Glycine (1.5 g/L) three times for 5 minutes each. Organoids were then blocked and permeabilized using a solution containing 10% donkey normal serum (Sigma), 0.5% Triton X-100 (Sigma), and 0.5% BSA in PBS on a thermal mixer at 300 RPM at 4 °C overnight. Organoids were then washed with PBS and incubated with primary antibodies in a solution containing 1% Donkey Normal Serum, 0.5% Triton X-100, and 0.5% BSA in PBS on a thermal mixer at 300rpm at 4 °C for 24 hours. Following this, organoids were washed with PBS. Organoids were then incubated with secondary antibodies on a thermal mixer at 300rpm at 4 °C for 24 hours in the dark. Subsequently, organoids were washed with PBS and mounted on glass microscope slides. To retain three-dimensionality, 90-μm polybead microspheres (Polyscience, Inc.) were placed between the slide and a No. 1.5 coverslip to provide support pillars. Organoids were imaged using a Nikon A1 Confocal. Images were analyzed using ImageJ. When comparing images across or between conditions, for each channel of an image being measured, pixel intensity values were equalized to that of the H1 WT condition. The circularity index was calculated using the following formula: Circularity Index = 4*pi*(Area) / (Perimeter^2).

### Measuring of cytosolic Ca^2+^ fluxes

Calcium imaging was performed as previously described^91^ with adaptation. In brief, cells were plated on 35 mm collagen-coated glass bottom dishes (MatTek). Cell culture media was removed, and cells washed once with tyrode’s buffer (140 nmM NaCl, 5.4 mM KCl, 10 mM glucose, 1 mM MgCl_2_, 2 mM pyruvate, 2 mM CaCl_2_, 10 mM HEPES). Cells were then loaded with Fura2-AM (Thermo Fisher) diluted to 2 µM in tyrode’s buffer for 30 minutes at 37°C. Cells were washed 2x in tyrode’s buffer before being imaged on a Carl Zeiss Axio Observer Z1 microscope. Traces were recorded in Zen Softward (Zeiss) and analysed using clampfit 10.7 (Molecular Devices).

### ChIP-qPCR and ChIP-seq procedures

ChIP experiments were performed as previously described^92^. In short, the cells were double crosslinked with 2mM di (N-succinimidyl) glutarate (DSG) for 45 minutes followed by 1% formaldehyde for an additional 15 minutes, at room temperature. Fixed cells were lysed and sonicated using a probe sonicator. The chromatin was then centrifuged at max speed for 30min. Supernatant containing the chromatin was collected and subjected to overnight incubation with specific antibodies at 4°C (**Table S6**). Subsequently, DNA purification was carried out using the Qiaquick PCR purification kit (QIAGEN, 28106). For ChIP-qPCR analysis, DNA was eluted in 75ul of elution buffer, with 2ul used for each qPCR reaction. Amplification was performed utilizing SYBR Green master mix and conducted in a QuantStudio3 instrument. ChIP-seq experiments were performed in the same fashion as ChIP-qPCR. The NEBNext^®^ Ultra™ II DNA Library Prep Kit was used for library preparation followed high-throughput sequencing using NextSeq1000.

### Generation of epiblast-specific conditional YAP1 knockout mice

The YAP^fl/fl^ and the Sox2cre mice were purchased from Jackson laboratories. Female Yap^fl/fl^ were crossed with epiblast specific Cre transgenic male mice, Sox2cre:Yap^fl/+^, to generate conditional YAP1 knockouts in the Sox2 lineage (Sox2cre:Yap^fl/fl^). Only males were carrying the Sox2cre allele were used for the crossing due to known maternal inheritance of this Cre^93^. All animal work complied with ethical regulations for animal testing and research and was done in accordance with IACUC approval by Temple University and followed all AAALAC guidelines. Mice were genotyped using standard flox and cre protocols (**Table S6**).

### Embryo isolation and scRNAseq procedure of E7.75 embryos

We followed procedures for the single-cell analysis of perigastrulating mouse embryos that we previously optimized in the Estaras lab^94^. To analyze conditional *Yap1* KO embryos, timed pregnancies were calculated from the noon of the day of the vaginal plug, considered E0.5. Thus, embryos were isolated seven days later at 3pm, which corresponds to E7.75. The visceral yolk sac was collected for same-day genotyping for Cre and flox. For the single-cell RNAseq experiments, multiple pregnancies were synchronized to obtain enough number of embryos with desired genotype. For the single-cell analysis, 10 cYap1 KO embryos (sox2cre:Yap^fl/fl^) and 7 heterozygous controls (sox2cre:Yap^fl/+^) were processed. After genotyping, embryos were pooled and dissociated using trypsin for 5 minutes at 37°C. After 5 minutes, trypsin was neutralized with three times the amount of DMEM/10% FBS. The 10x Chromium Next GEM Single Cell 3’ GEM, Library & Gel Bead Kit v3.1 was used. Generation of gel beads in emulsion, barcoding, GEM-RT clean-up, complementary DNA amplification, and library construction were all performed as per the manufacturer’s protocol. Qubit and tapestation were used for library quantification before pooling. Libraries were pooled and sequenced using NextSeq2000.

### scRNAseq analysis of E7.75 embryos

Single cell sequencing reads were counted individually for each sample using cellranger (v7.1.0) with default parameter. Detection of doublet and cells contaminated with ambient RNA were assessed using scrublet and soupX (v1.6.2), in R and Python, respectively. Low-quality cells, including doublets, cells with over 25% mitochondrial content, and those expressing fewer than 2000 RNA features, were filtered out. Most analysis were performed using the Seurat R package (v4.3.0), including quality control plots generated with FeatureScatter and VlnPlot Seurat’ functions. Samples were normalized and variance stabilized using the SCTransform function in Seurat, regressing out variables including nCount_RNA, percent.mt, percent.rb, S.Score, and G2M.Score. Principal component analysis (PCA) was run with 19 principal components for both samples. The samples were then integrated using Seurat’s SelectIntegrationFeatures, PrepSCTIntegration, FindIntegrationAnchors, and IntegrateData functions with 3000 selected features and the SCTransform normalization method. For visualization of gene expression in UMAP, and heatmap, the SCT, and RNA assay were used, respectively. Differential gene expression analysis for each cluster between genotypes was performed on the RNA assay using the FindMarkers Seurat function after log normalization and scaling. Genes were considered differentially expressed when adjusted p-value, based on Bonferroni correction was below 0.05. Cell type count differences between genotypes for each cell types were assessed by randomly downsampling WT samples 100 times to the same number of cells as the cYap1 KO sample (3,621 cells). For each iteration, the number of cells in each cluster was counted. Chi-squared tests with Bonferroni correction were used to assessed significant differences in cell counts between genotypes across clusters. Pathway analysis was performed using the SCPA R package (v1.5.4) with curated gene lists selected from the MSigDB C2 database. Pseudotime trajectory analysis was generated using the condiments R package (v1.2.0). Trajectory inference was performed with the Slingshot package (v2.2.1) and visualized with the ggplot2 R packages. Gene ontology analysis was performed using the enrichR R package (v3.2) using the GO_Biological_Process_2023 library and visualized with ggplot2.

### scATACseq of hESC-CPCs and CMs and bioinformatics analysis

WT and YAP KO H1 Day 5 cell cultures (untreated and RA-treated) and Day 30 CMs (RA-treated) were processed for single-cell RNAseq following procedures shown in **Figure 1A**. Two differentiation experiments were pooled for gem generation and library preparation using the 10x Chromium Next GEM Single Cell 3’ GEM. Single cell sequencing reads were aligned to human reference genome hg19 and then counted individually for each sample using cellranger (v2.1.1) with default parameter. For quality-control filtering, aligned cell and transcript counts from each sample were processed by Seurat R package (v3.9.9) by removing low-quality cells and doublets containing over 10% of mitochondrial reads, fewer than 200 detected genes and over 10,000 detected genes. Samples were log normalized with NormalizedData function and the NormalizeData function and the top2K variable genes were identified with the FindVariableFeatures function. Principal component analysis (PCA) was run with 10 principal components for both samples. The samples were then integrated using Seurat’s merge function. Clusters were identified with the FindClusters function with a clustering resolution set to 0.4 and 0.8 for two datasets, respectively. Clusters with only one cell were removed. Differential gene expression analysis for each cluster between genotypes was performed on the RNA assay using the FindMarkers Seurat function with default Wilcoxon Rank Sum test. Genes were considered differentially expressed when adjusted p-value, based on Bonferroni correction was below 0.01 and absolute value of fold change in log2 scale was higher than 0.5.

### Single-nuclei ATAC-sequencing (snATAC-seq) of hESC-CPCs and bioinformatics analysis

WT and YAP KO H1 Day 5 cell cultures (untreated and RA-treated) were used for snATAC-seq analysis. Two differentiation experiments were pooled for downstream processing. Sample preparation, library generation and single-cell sequencing snATAC-seq were carried out at the Center for Epigenomics at UCSD. The procedure followed the protocol described in Cusanovich et al(Cusanovich et al., 2015). Briefly, DMSO-frozen hESCs were thawed in 37°C water bath until liquid (∼2-3 minutes) and immediately placed on ice. Nuclei were isolated and permeabilized using OMNI buffer (0.1% NP40, 0.1% Tween-20, 0.01% Digitonin) as described(Corces et al., 2017). Nuclei were counted, and 2k nuclei were dispensed per well; 96 wells per sample for tagmentation. After tagmentation (and addition of the first barcode), nuclei from individual samples were pooled. 20 nuclei from each sample were sorted per well (768 wells total). Following nuclear lysis, the second barcode was added, and DNA molecules were pooled and purified. Before sequencing, the quality of the library was certified by analyzing fragment size distribution and DNA concentration. Sequencing was performed in a HiSeq 4000 device.

SnapTools (v1.4.7) and SnapATAC (v1.0.0) developed at Ren’s Lab, UCSD were used for the scATAC-seq analysis(Fang et al., 2021). In brief, reads were map ped to hg19 reference using bwa mem (v0.7.12)(Li and Durbin, 2009) and sorted by samtools (v1.9)(Li et al., 2009). Then reads were pre-processed by SnapTools using default parameters and chromosome bin size of 5000. Cells with less than 500 unique molecular identifier (UMI) reads were removed from downstream analysis. The distribution of counts in promoter ratio was inspected, and cells with the ratio < 0.1 or > 0.6 were removed. MACS2 (v2.1.2)(Zhang et al., 2008) was used for peak calling from pooled reads (parameters: -g hs --nomodel --shift 37 --ext 73 - -qval 1e-2 -B -- SPMR --call-summits). Chromosomal bins overlapped with the ENCODE hg19 blacklist regions, random genomic contigs, or mitochondrial chromosomes were removed from the analysis. Top 5% of the most expressed bins were also removed. Dimension reduction by Latent semantic analysis (LSA) was performed on binary chromosomal bins and visualized by UMAP using the top 30 principal components and a random seed of 10. Differential regulated (DR) regions were called by SnapATAC command “findDAR” using parameters “bcv=0.1, fdr=0.05, pvalue=0.01, test.method=exactTest, seed.use=10”. DR regions with false discovery rate (FDR) < 0.05 and absolute log2 fold-change > 1 were considered as significantly different. Motifs enrichment analysis was performed by HOMER (v4.9.1)( http://homer.ucsd.edu/homer/), with parameters “-size 200 -mask”. The nearby genes associated with DR regions were annotated with ‘annotatePeaks’ program from HOMER with default parameters. Gene ontology analyses of the associated genes were carried out by over-representation analysis (ORA) using WebGestaltR R package. Pathways with FDR<0.05 were considered as significantly enriched and were visualized with enrichment ratio. To calculate the distribution of normalized read coverage round the center of DR regions, HOMER ‘annotatePeaks’ program was used with a window size of +/-3kb and a bin size of 25bp. The normalized read coverage was visualized in heatmap.

### ChIP-seq analysis

ChIP fragments were sequenced in an Illumina sequencer. Reads were aligned to the Human hg19 genome assembly (NCBI Build 37) using STAR (v2.5.1b, doi: 10.1093/bioinformatics/bts635. pmid:23104886) with default parameters. Reads soft-clipping and splicing were turned off by specifying ‘--alignEndsType EndtoEnd -- alignIntronMax 1’ for ChIP-Seq mapping. Only reads that mapped uniquely to the genome were considered for further analysis. Peak finding, motif finding and peak annotation, genome browser read density files were performed using HOMER (v4.9.1). Peaks were identified using ‘findpeaks’ program, using input condition as background reference and default parameters (four-fold enrichment over input control, four-fold enrichment over local tag count, a Possion p-value of 1e-4, and style factor). Peaks were annotated by associating with the closest RefSeq defined TSS by the ‘annotatePeaks’ program in HOMER. Overlapping peaks of YAP ChIP-seq with RA-ATACseq peaks was performed using ‘mergePeaks’ with parameters ‘-d given’.

### Data accessibility

Th genomic sequencing datasets were submitted to GEO database and it will be available after peer-reviewed publication.

## Supplementary Figures

**Figure S1.**
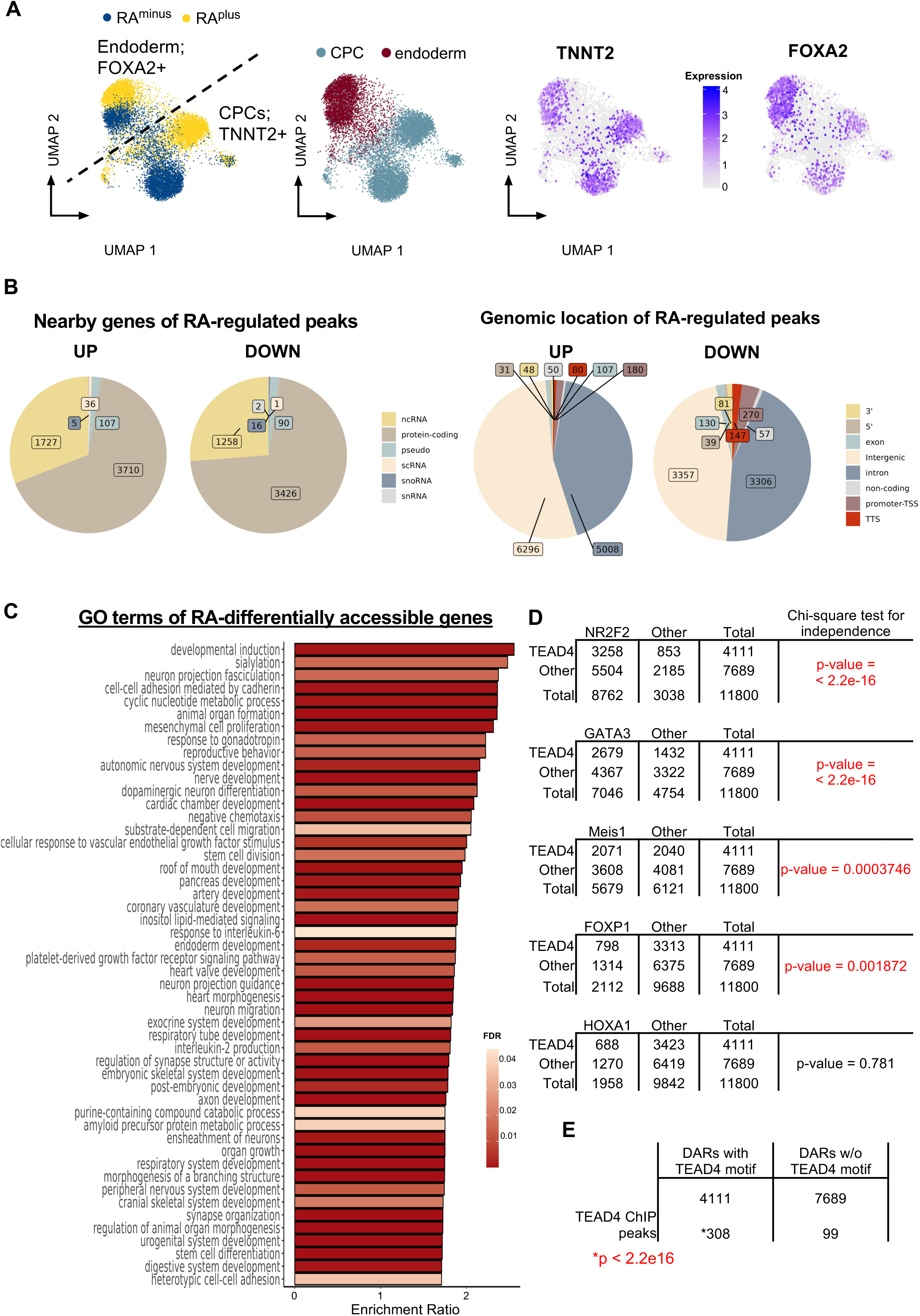
**A)** UMAP projection of snATACseq experiments in RA^minus^ and RA^plus^ hCPCs depicting two predominant populations (endoderm and CPCs). The UMAPs on the right show the accessibility for FOXA2 gene (endoderm marker) and TNNT2 (cardiac marker). **B)** Pie charts show distribution of RA-regulated peaks detected in the snATACseq experiments. **C)** Gene Ontology analysis of RA-differentially accessible genes. **D)** Bar graph showing the frequency of TEAD4 and other atrial transcription factors (NR2F2, GATA3, MESI1, FOXP1, and HOXA1) motifs in RA-induced sites. Note that HOXA1 was not significantly associated to TEAD4 enhancers. **E)** Table displays the number of TEAD4 peaks (ChIP-seq) associated to DARs with and without TEAD4 motifs in RA-treated hESC-CPCs. Chi-square test for independence.

**Figure S2.**
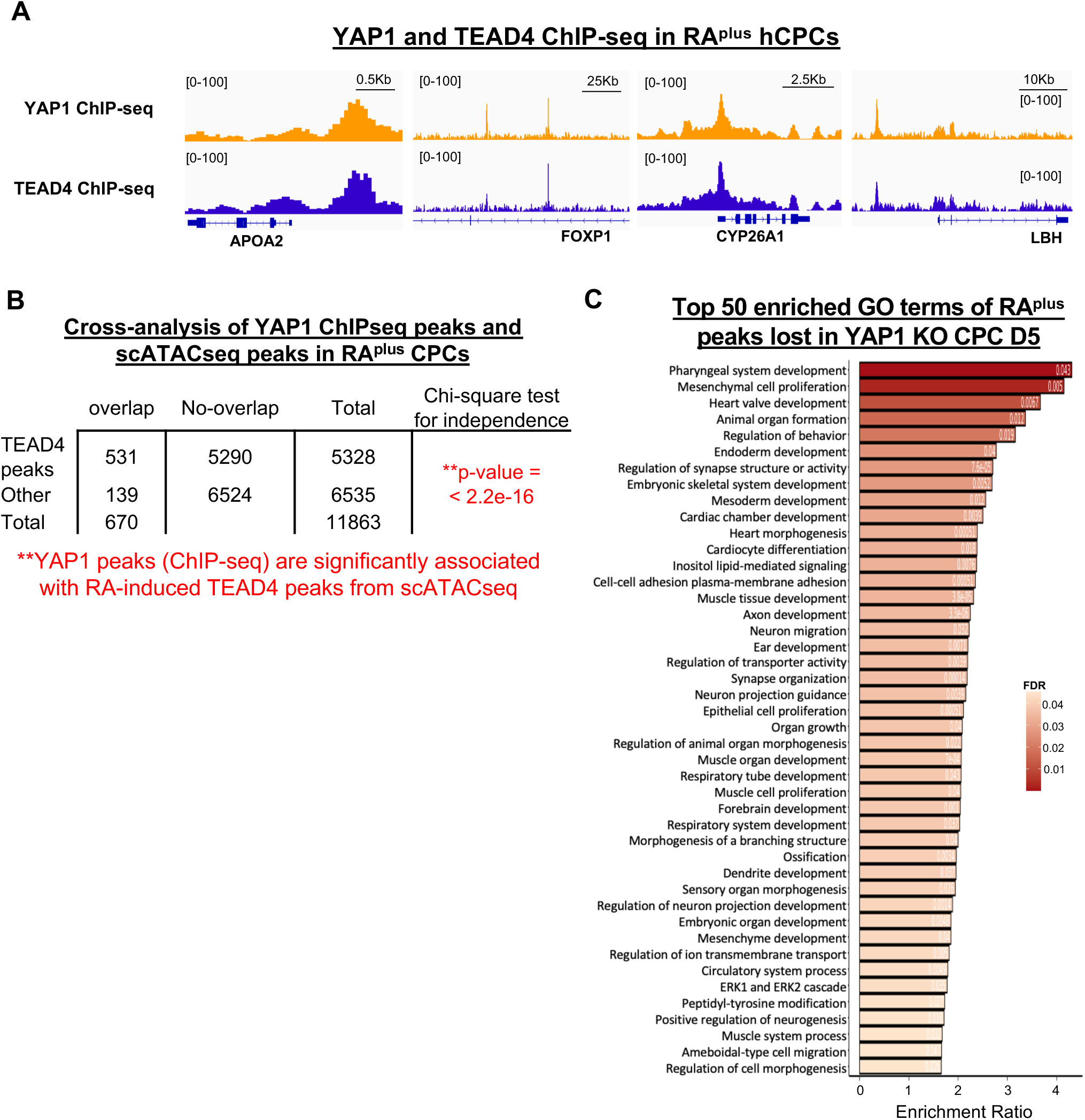
**A)** Genome browser captures show YAP1 and TEAD4 binding peaks on indicated genes, in RA^plus^ hCPCs. **B)** Table shows association between YAP1 binding peaks (ChIP-seq) and RA-induced TEAD4 enhancers (snATACseq) with corresponding statistical analysis. **C)** Gene Ontology analysis of RA-induced chromatin regions that lost accessibility in the YAP1 KO RA^plus^ hCPCs.

**Figure S3.**
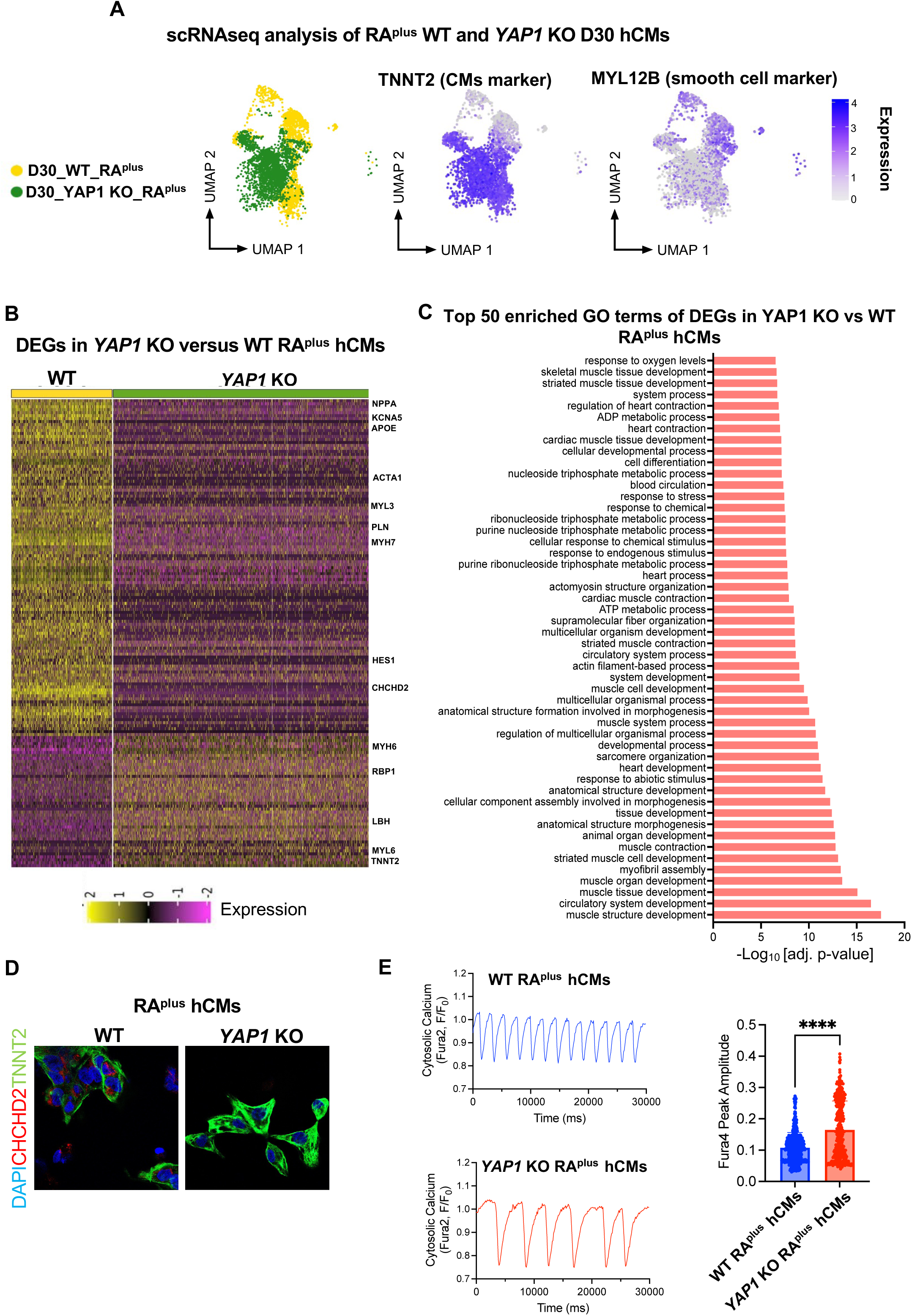
**A)** UMAP distribution of WT and YAP1 KO Day 30 cultures derived from RA-treated hCPCs. Two main populations were identified corresponding to CMs (TNNT2+) and smooth muscle cells (MYL12B+). **B)** Heatmap shows differentially expressed genes in WT and YAP1 KO RA^plus^ hCMs (abs (Log2FC)>0.5, adj p <0.05). **C)** GO terms enriched in the DEGs identified in the YAP1 KO vs WT RA^plus^ hCMs. **D)** Representative immunostaining of indicated markers in WT and YAP1 KO RA^plus^ hCMs. **E)** WT and YAP1 KO RA^plus^ hCMs were stained with the Fura4 cytosolic calcium and peak amplitude were measured and quantified. Representative traces and peak amplitude are shown on the right. n=30-40 cells/group and images:8-15/group.

**Figure S4.**
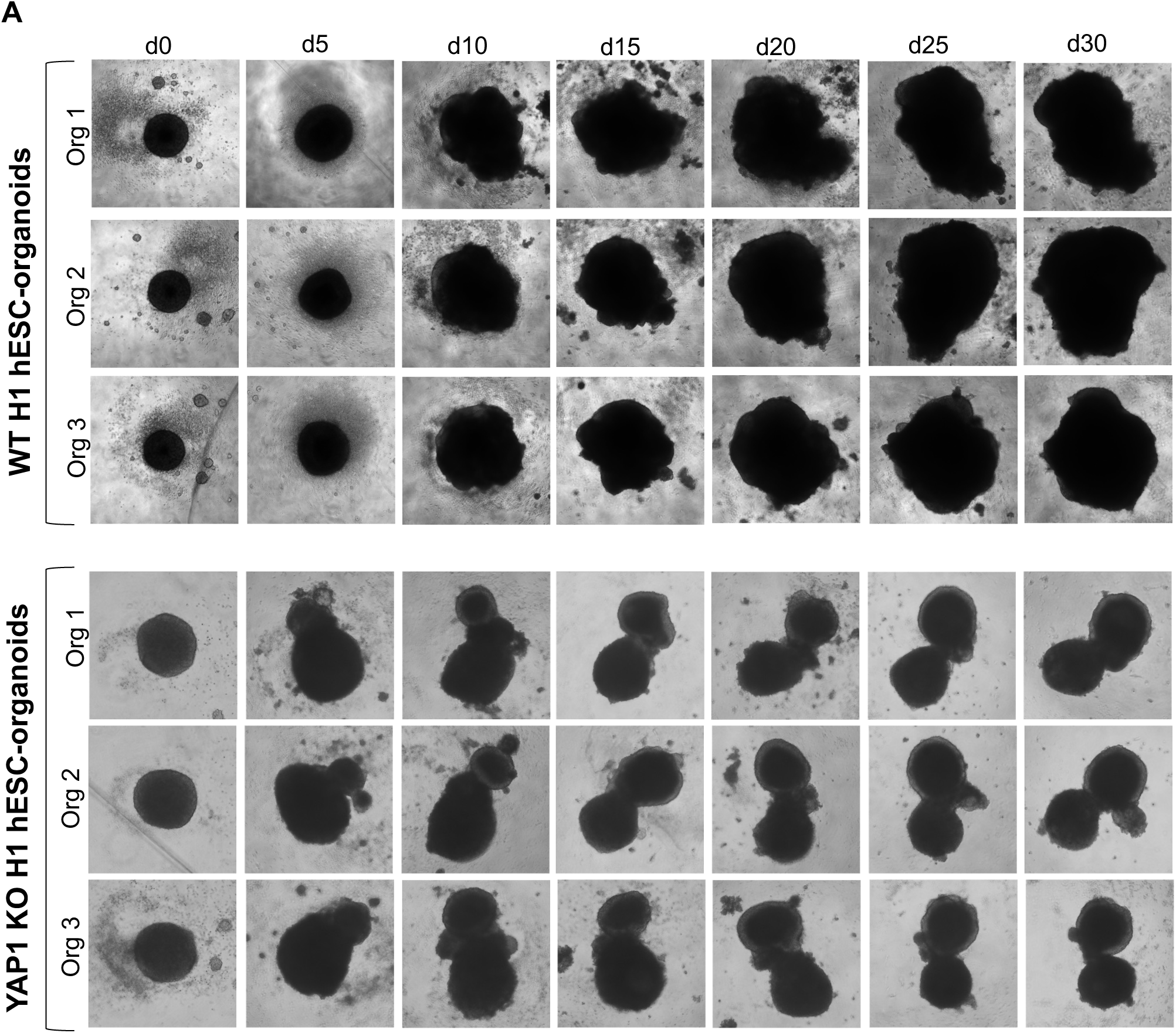
**A)** WT and YAP1 KO H1 hESCs were differentiated to cardiac organoids and bright field images were taken at the indicated days. Three organoids per condition are shown.

**Figure S5.**
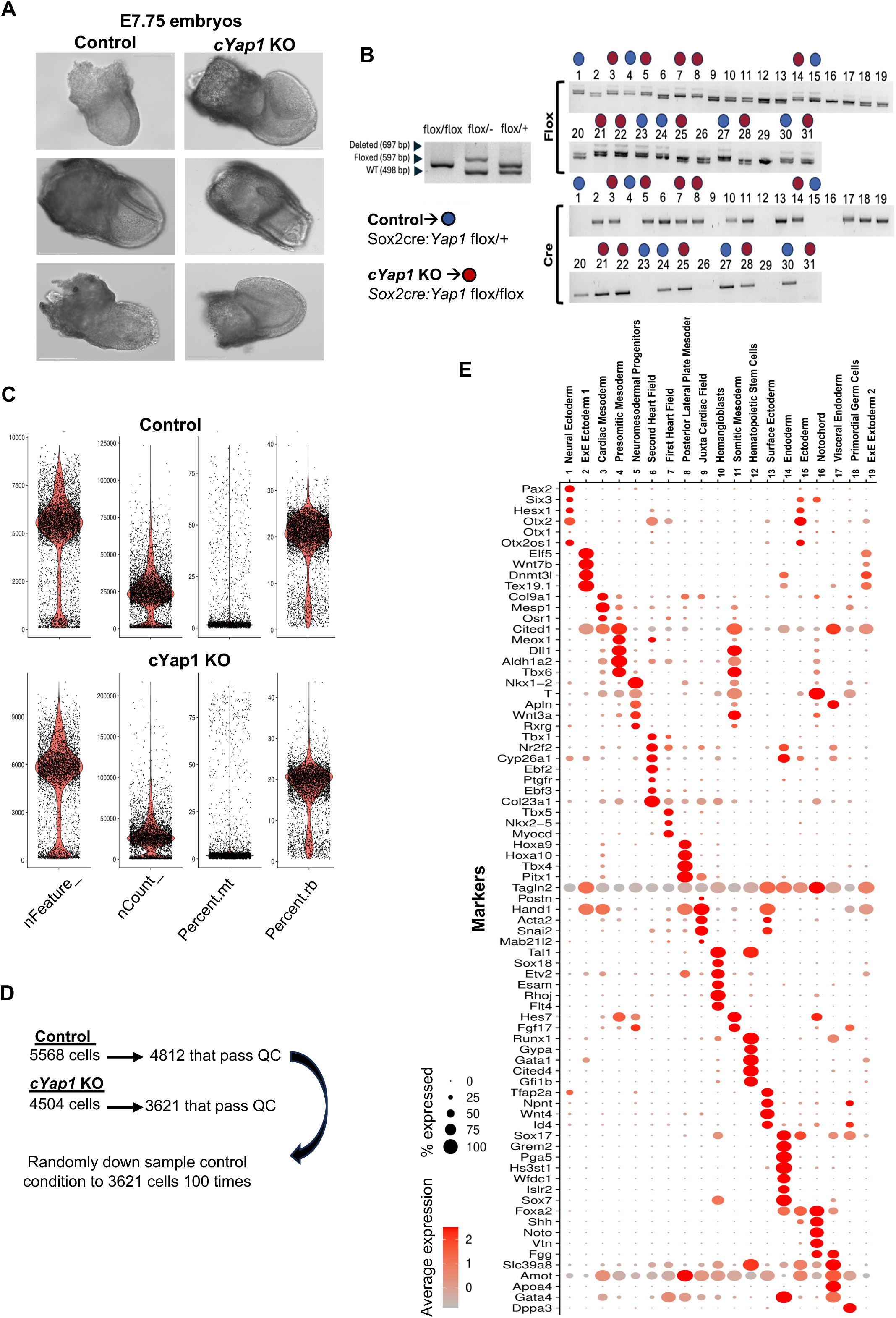
**A)** Representative images of three E7.75 control and *cYap1* KO embryos. **B)** Genotyping results for cre and flox alleles of the embryos processed for scRNAseq analysis. **C)** Graphs showing quality controls parameters of scRNAseq of control and *cYap1* KO embryos. **D)** Total number of control and *cYap1* KO cells sequenced that passed the QC. The downstream analysis was performed with 100 bootstrap iterations for downsampling wild-type (WT) cells to 3,621 cells each time. **E)** Dot plot shows the expression levels of the markers used to annotate each of the 19 cell populations.

**Figure S6.**
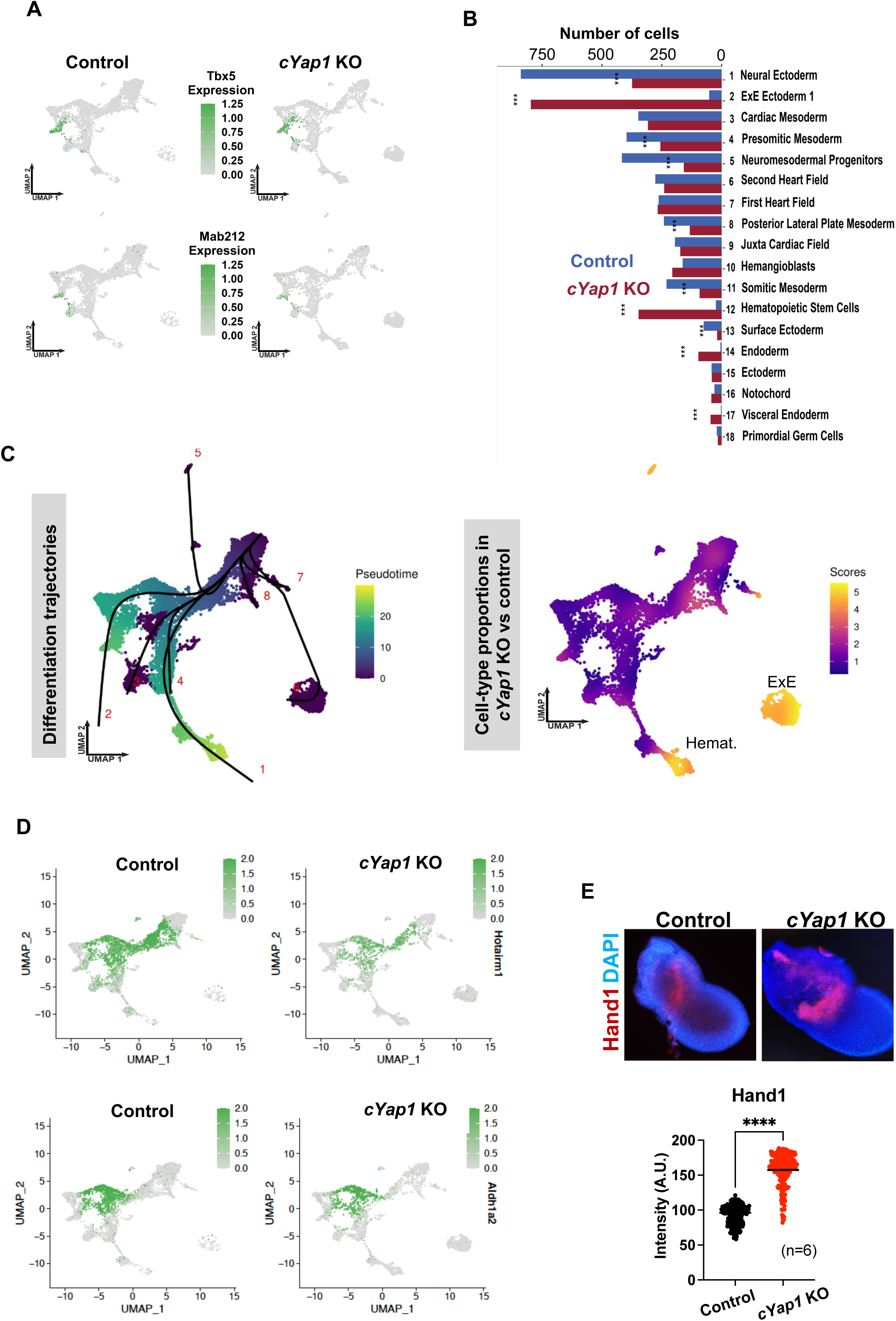
**A)** UMAP showing *Tbx5* (FHF marker) and *Mabl2* (JCF marker) expression levels in control and *cYap1* KO embryos. **B)** Bar graph indicates the normalized number of cells in each cluster in control and *cYap1* KO embryos. Stars indicate statistical differences determined by a chi-squared test with a significance threshold of <0.05 (*), <0.001 (**), and <0.0001 (***). The analysis was performed with downsampling wild-type (WT) cells to 3,621 cells each time. P-values were adjusted using the Bonferroni correction method. **C)** UMAP displaying the 8 cell trajectories colored by pseudotime (left). UMAP highlighting the difference in cell type proportions between WT and *cYap1* KO (right; the higher the score, the greater the difference between genotypes). Note that the extraembryonic trajectory (6) and the hematopoietic trajectory (1) were accelerated in the *cYap1* KO embryos. **D)** UMAPs showing *Hotairm1* and *Aldh1a2* expression levels in control and *cYap1* KO embryos. *Hotairm1* was significantly downregulated in 9 of the 19 clusters (LogFC>0 adj p<0.05). A downregulation trend is observed in *Aldh1a2* in several clusters in the *cYap1* KO embryos, including in the SHF (Log2FC= -1.14, p value=0.000398, Adj p value, n.s). **E)** RNAscope analysis of *Hand1* probe in E7.5 control and *cYap1* KO embryos. Graph shows quantification of fluorescence signal (n=5 embryos, p<0.0001, t-test). Note that scRNAseq also detected upregulation of *Hand1* in the nascent mesoderm cluster of the *cYap1* KO embryos, compared to controls (Log2FC= 0.43, Adj p value 0.00014).

## Author contributions

C.E. conceived the project, performed hESCs-differentiation experiments, supervised and designed experiments, and attained funding.

E.A. performed hESC-experiments, ChIP-qPCRs, library preps and the experiments on mouse embryos.

C.E and E.A made figures and wrote the manuscript.

B.V and A.A performed cardiac organoid experiments and analysis.

L.H, J.Y and A.W performed all bioinformatics analysis of the in vitro data, generated the corresponding figures, and edit manuscript.

E.S performed the HAND1 WBs and help with the animal experiments.

T.R and N.A analyzed the single-cell RNAseq of the embryos, make corresponding figures and edit manuscript.

H.C and J.E performed the calcium analysis of the hESC-CMs.

E.M and E.A performed ChIP-qPCR experiments on the CPCs.

A.D and E.A performed RNAscope experiments and quantification.

R.F performed initial analysis of the ChIP-seq experiments.

A.M performed initial analysis on scRNAseq of embryos.

## Acknowledgements

We acknowledge the Whetstine lab (Dr. Zach Grey, Madison Honer, Benjamin Ferman, and Dr. Johnathan Whetstine) and the Genomics Facility at the Fox Chase Cancer Center for technical support for single-cell experiments. We acknowledge support from Salk Institute Bioinformatics and Stem Cell Cores, and the center for Epigenomics at the University of California, San Diego (UCSD).This work is funded by the NIH/NHLBI R56HL163146 (to C.E), NIH/NHLBI T32 training grant 5T32HL091804 to E.A, W.W. Smith Charitable Trust grant (to C.E.), and the California Institute for Regenerative Medicine (to Kathy Jones, former C.E advisor, GC1R-06673-B).

